# TyCHE enables time-resolved lineage tracing of heterogeneously-evolving populations

**DOI:** 10.1101/2025.10.21.683591

**Authors:** Jessie J. Fielding, Sherry Wu, Hunter J. Melton, Crystal Z. Wang, Nic Fisk, Louis du Plessis, Kenneth B. Hoehn

## Abstract

Phylogenetic methods for cell lineage tracing have driven significant insights into organismal development, immune responses, and tumor evolution. While most methods estimate mutation trees, time-resolved lineage trees are more interpretable and could relate events like cellular migration and differentiation to perturbations like vaccines and drug treatments. However, somatic mutation rates vary dramatically by cell type, significantly biasing existing methods. We introduce TyCHE (Type-linked Clocks for Heterogeneous Evolution), a Bayesian phylogenetics package that infers time-resolved phylogenies of populations with distinct evolutionary rates. We demonstrate that TyCHE improves tree accuracy using a new simulation package SimBLE (Simulator of B cell Lineage Evolution). We use TyCHE to infer patterns of memory B cell differentiation during HIV infection, dynamics of recall germinal centers following influenza vaccination, evolution of a glioma tumor lineage, and progression of a bacterial lung infection. TyCHE and SimBLE are available as open-source software packages compatible with the BEAST2 and Immcantation ecosystems.

## Introduction

Human diseases are fundamentally shaped by cell proliferation, differentiation into subtypes, and migration across tissues. Phylogenetic methods for cellular lineage tracing have driven significant recent insights into development and disease.^1–7^ These methods either use mutations from implanted dynamic lineage recorders or naturally-occurring somatic mutations.^8–11^ While most phylogenetic methods estimate genetic distance trees in which branch lengths represent mutations per site, some approaches estimate time trees in which branch lengths represent calendar time.^12^ Time trees have significant advantages because they provide calendar date estimates for evolutionary events and can therefore relate them to external perturbations. For example, they have been used to identify the ignition date of the HIV-1 pandemic^13,14^ and the spread of SARS-CoV-2 globally.^15,16^ Time trees also contain a signature of past population dynamics^17^ and can be used to infer epidemiological parameters like the migration history and changes in effective population size, for example quantifying the effect of containment measures during recent Ebola virus and SARS-CoV-2 epidemics.^18–20^ Applied to cell populations, time trees could be used to infer cell migration, differentiation, and proliferation in response to perturbations like vaccines, antibiotics, and chemotherapy.^21,22^

A major challenge for inferring time trees for cell populations is that cellular evolution is highly heterogeneous. While many viruses evolve in a clock-like manner, somatic mutations in cells are often linked to “hypermutator” states. For example, B cells undergo periods of rapid somatic hypermutation during immune responses before becoming quiescent memory cells.^23^ Infectious bacterial populations, viruses, and tumors also frequently develop hypermutator phenotypes.^24–27^ Even artificial, dynamic lineage recorders are frequently designed to mutate only during distinct periods induced by, for example, doxycycline.^28,29^ Existing methods for estimating time trees typically assume the evolutionary rate is uncorrelated with cellular phenotypes, leading to biased inferences when applied to cell populations with very different evolutionary rates.^12,30^

Here, we introduce TyCHE (Type-linked Clocks for Heterogeneous Evolution), which integrates with the BEAST2^31^ Bayesian phylogenetics platform and estimates accurate time trees of heterogeneously evolving populations. In contrast to existing methods, TyCHE simultaneously reconstructs ancestral cell type states and estimates the dates of tree nodes using separate molecular clock rates for each cell type. To validate TyCHE, we also developed SimBLE (Simulator of B cell Lineage Evolution) which simulates heterogeneously evolving populations, including realistic B cell germinal center (GC) reactions. Using simulations, we demonstrate that TyCHE estimates time trees with more accurate dates, topologies, and reconstructed cell types than existing methods. Using empirical B cell receptor (BCR) sequences, we show that TyCHE can infer realistic patterns of B cell evolution and memory B cell formation during chronic HIV infection^32^ as well as realistic dynamics of recall GCs following repeated influenza vaccination.^33^ Further, we use TyCHE to reconstruct the temporal evolution of a hypermutating glioma tumor lineage, as well as the evolution of a *Pseudomonas aeruginosa* hypermutator lineage during an acute lung infection.^25,34^ TyCHE and SimBLE both interface with the R package Dowser for cellular phylogenetic analyses, which is part of the Immcantation suite.^35^

## Results

### TyCHE: Type-linked clock models for heterogeneous evolution

Inferring time-resolved phylogenies requires a clock model, which describes the relationship between evolution and time. A “strict clock” (SC) assumes a constant rate of evolution. Other models, such as the uncorrelated lognormally distributed (UCLD) relaxed clock, allow rates to vary.^30,36^ Bayesian time tree estimation typically uses Markov Chain Monte Carlo (MCMC), in which tree topologies, node heights (dates), and model parameters are iteratively proposed. For each proposal, the clock model converts branch lengths from calendar time to genetic distance, enabling computation of the sequence data likelihood.^37^ New topologies, node heights, and parameter values are accepted or rejected based on their posterior probabilities relative to the current state.^12,31^

In contrast to prior approaches that model trait values as independent of the clock rate, TyCHE links each cell type to a separate molecular clock rate which can be either fixed or estimated as a parameter (**Fig. 1**). These clock rates can be *a priori* estimated using root-to-tip regression or by fitting a strict clock model to pure cell populations.^12,38^ Each node is assigned a cell type and the genetic distance of each branch is calculated based on its length and the states of its parent and child nodes. The cell types at each node are modeled as a continuous time Markov chain (CTMCs). This CTMC is parameterized by a transition rate matrix, which can be used to estimate the relative rates of cellular transitions and can also incorporate prior information such as irreversibility of cell differentiation events. We further use these relative rates to calculate the expected occupancy time in each cell type between nodes. Critically, in contrast to prior approaches that treat trait values as conditionally independent of the sequence data likelihood, TyCHE integrates across cell type states at internal nodes using MCMC.^39^

**Figure 1:**
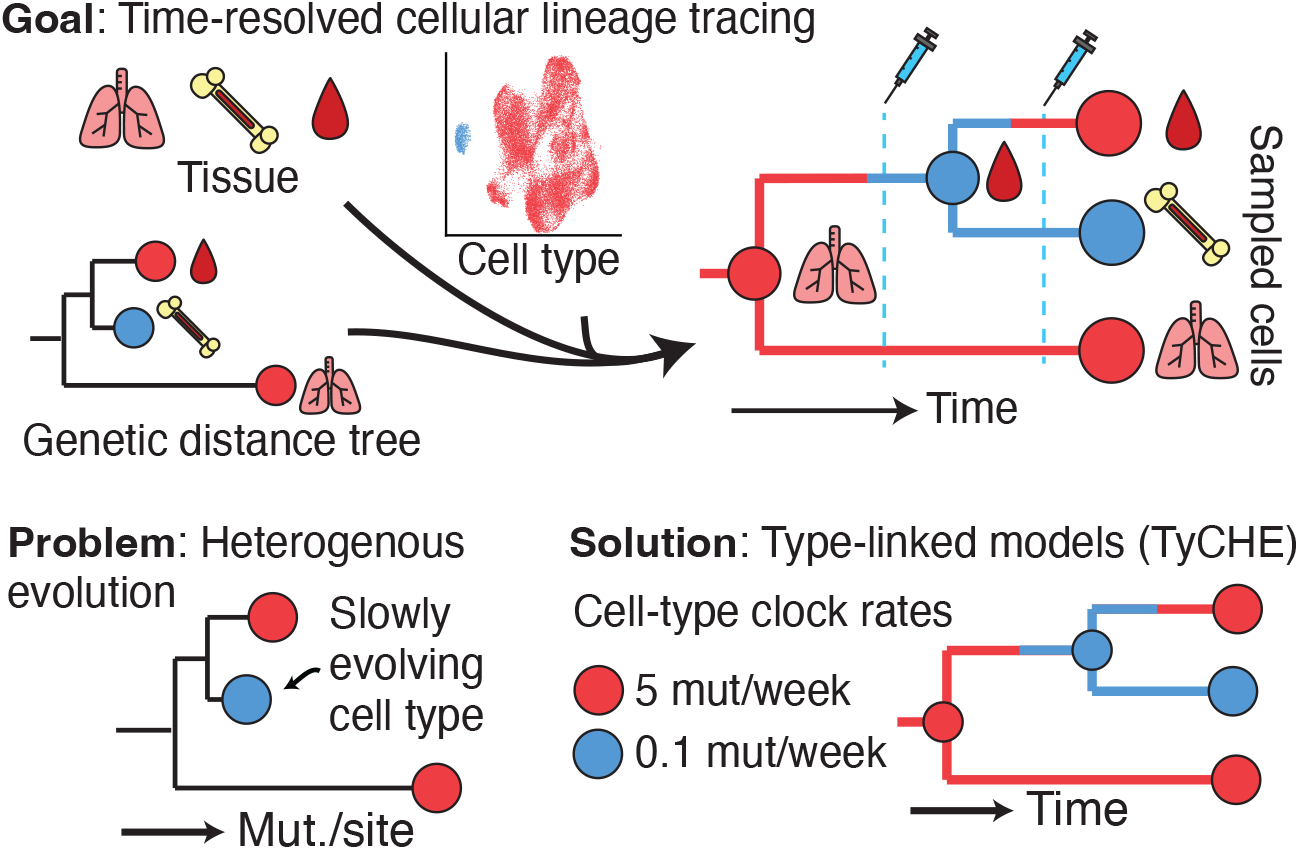
Graphical representation of TyCHE. Time-resolved cellular lineage tracing has the potential to unravel the dynamics of cellular evolution, migration, and differentiation. However, existing molecular clock models are inappropriate when evolutionary rates are linked to discrete cell types. TyCHE solves this problem using type-linked clock models that allow cell types to evolve at distinct rates.

### SimBLE: Simulating germinal center reactions

B cell evolution is highly heterogeneous. B cells produce antibodies, whose structures are also expressed as membrane-bound B cell receptors (BCRs). During immune responses, B cells can enter germinal center (GC) reactions, during which they upregulate enzymes that rapidly introduce mutations in a process called somatic hypermutation (SHM).^23^ GC B cells with affinity-improving mutations survive and proliferate, while others undergo apoptosis. B cells can differentiate into memory B cells (MBCs) which can be stimulated upon re-infection, or plasma cells which migrate to the bone marrow and release antibodies. In contrast to GC B cells, neither MBCs nor plasma cells undergo SHM, so time-resolved phylogenetic analyses of B cells must take these vastly different evolutionary rates into account.

To benchmark TyCHE, we developed SimBLE, an agent-based simulator of heterogeneous evolution, including B cell evolution and differentiation in GCs (**Fig. 2A**). Briefly, a starting pair of BCR heavy and light chain sequences is randomly chosen from a dataset of naive single B cells.^35,40^ GC B cells mutate according to models of SHM targeting, and affinity is calculated based on similarity to a target amino acid sequence.^41,42^ B cells proliferate proportionally to their relative affinity. GC B cells differentiate into MBCs early on before shifting to mainly plasma cell production.^43^ SimBLE incorporates recent discoveries, including silencing of SHM during clonal bursts and a log-additive relationship between mutations and affinity.^44–46^

**Figure 2:**
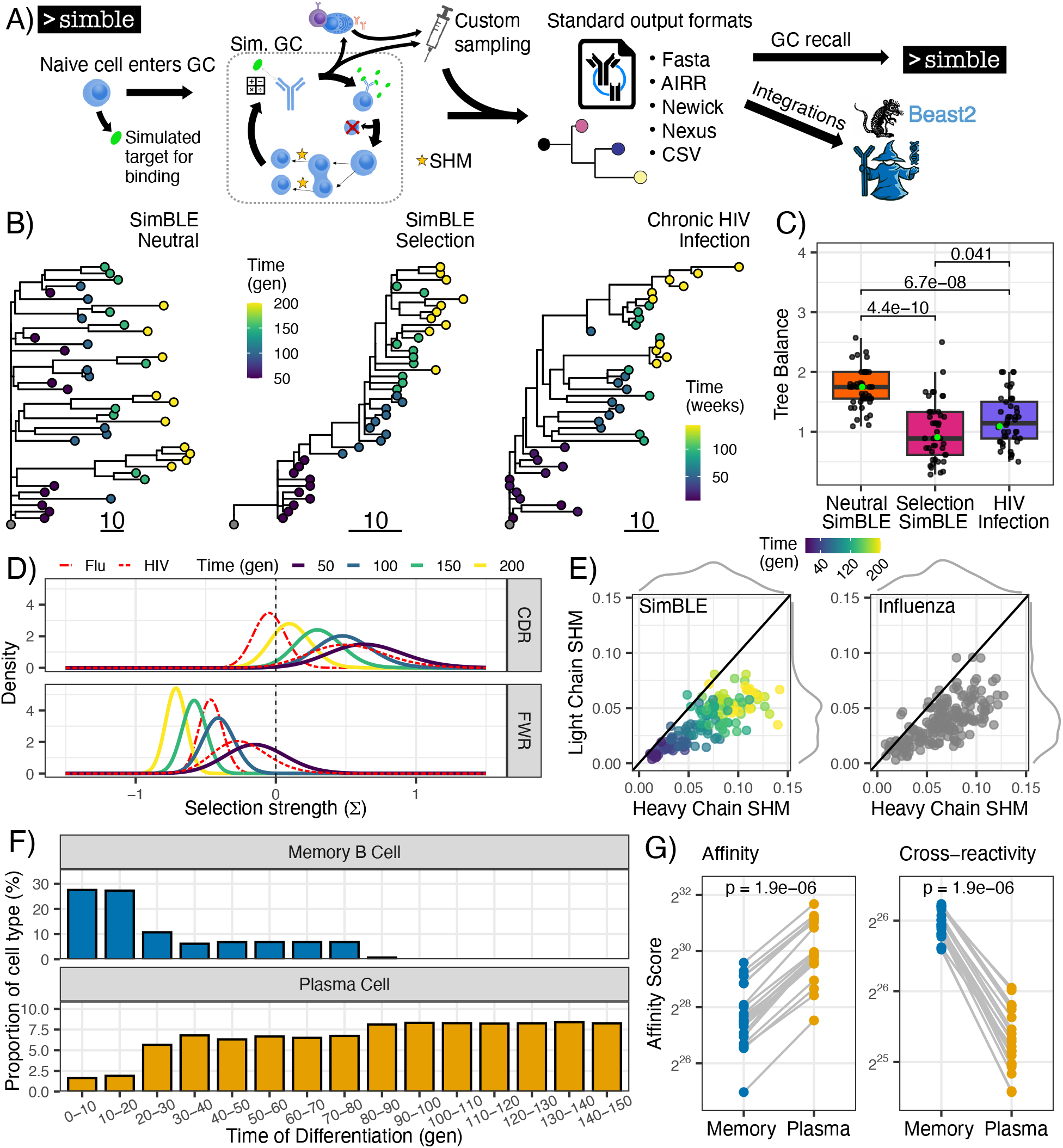
Simulating GC reactions and other type-linked evolution using SimBLE. **A)** Graphical representation of the SimBLE algorithm. **B)** Estimated genetic distance trees from BCR sequences simulated under neutral and selective evolution, as well as previously published BCRs from chronic HIV infection.^47^ All lineages were downsampled to 40 sequences. Tips are colored by sample time in generations (gen) or weeks. **C)** Tree balance measured as maximum width over maximum depth^53,54^ for simulations of neutral and selective evolution for 200 generations, and published BCRs from measurably evolving lineages from an HIV-1 infected donor.^47^ Each dataset used 45 lineages downsampled to 40 sequences each. Trees from panel B are highlighted in green. **D)** Posterior distributions of selection scores from BASELINe applied to BCRs from 100 lineages sampled at specified timepoints under simulations of selection. Also shown, in red, are posterior distributions of selections scores from BASELINe applied to the chronic HIV BCRs used in panel C and the post-influenza vaccination BCRs used in panel E. Values less than zero indicate purifying selection while values greater than zero indicate diversifying selection. **E)** BCR heavy and light chain mutation frequency. Fifteen lineages were simulated under selection for each specified number of generations for a total of 150 lineages. Each dot represents the mean of a lineage. Marginal distributions are shown on each axis. Line shows equal SHM in both chains. Single cell BCRs sampled following influenza vaccination are also shown.^35,48^ **F)** Proportion of MBCs and plasma cells by time of differentiation from the GC in 100 simulated lineages with a migration rate of 5 cells/generation. **G)** Mean affinity to target sequence (Affinity) and mean affinity to 20 target sequences differing from the original by 5 heavy and 3 light chain AA substitutions (Cross-reactivity). Twenty lineages were simulated for 150 generations. Each dot shows the mean of that cell type in one lineage, with lines connecting cell types from the same lineage. P values were calculated using the Wilcoxon signed-rank test.

We compared BCRs simulated from SimBLE to empirical BCR sequences from chronic HIV infection and influenza vaccination.^47,48^ Trees simulated under selection had imbalanced topologies similar to B cell trees from chronic HIV infection, while trees simulated under neutral evolution were more balanced (**Fig. 2B-2C**). BCRs simulated under selection showed diversifying selection in antigen-binding regions and purifying selection in structural framework regions, similar to real BCRs (**Fig. 2D**).^49^ Selection shifted towards purifying selection over time, consistent with prior studies.^50–52^ These signatures of selection were absent in BCRs simulated under neutral evolution (**Supplemental Fig. 1**). Simulated BCRs had realistic SHM levels (5-10%) that were roughly twice as high in the heavy chain, consistent with prior studies (**Fig. 2E**).^35^ Site-wise substitution patterns were distinct among selection, neutral, and uniform neutral simulations (**Supplemental Fig. 2**). MBCs differentiated before plasma cells, closely matching prior estimates (**Fig. 2F**).^43^ Further, while plasma cells had higher affinity to their target sequence, MBCs showed higher affinity to similar non-target sequences, representing broader reactivity (**Fig. 2G**). These results confirm SimBLE produces realistic GC and non-GC BCR sequences.

### Simulation-based validation of TyCHE

We validated TyCHE’s performance using simulated sequences of primary and recall GCs from SimBLE. Specifically, we compared TyCHE to the existing SC and UCLD relaxed clock models.^22^ For TyCHE, the prior GC clock rate mean was estimated using the mean strict clock rate estimated after removing non-GC B cells in each simulation. The clock rate for non-GC B cells was set arbitrarily low at 10^−6^ SHM/site/generation. For SC and UCLD models the clock rate, tree heights, and other parameters were estimated using all sequences simultaneously, as is standard. In simulated primary GC reactions, GC to non-GC transitions were considered irreversible in all models. In simulated recall GC reactions, GC to non-GC transitions were reversible, but the root cell type was fixed to GC and the maximum tree height was set to 1000 generations (∼83%) above the true tree height. These constraints are biologically justifiable in practice, and initial testing showed that they substantially improve convergence and identifiability (**Supplemental Fig. 3**).

Each simulation scenario was repeated to produce 20 lineages from distinct naïve BCR sequences. In primary GC simulations (**Fig. 3A**), B cells were sampled at generations 50, 100, 150, and 200. In recall GC simulations (**Fig. 3B**), B cells were sampled at generations 50 and 100 during a primary GC reaction. This GC reaction stopped at generation 100, and an MBC was randomly selected to initiate a recall GC at generation 1100, from which B cells were sampled from at generations 1150 and 1200. Model performance was evaluated based on tree height, total tree length, and Robinson-Foulds distance from the true tree topology. Tree height in particular is important, as it represents the initiation date of each GC reaction. TyCHE significantly outperformed existing models in both primary and recall GC simulations (**Fig. 3C**). These results remained consistent when these simulations were repeated using neutral evolution (**Supplemental Fig. 4**). These results demonstrate that TyCHE significantly outperforms existing clock models when clock rates vary by cell type in both the continuous, irreversible evolution of primary GC reactions and the start/stop dynamics of recall GC reactions.

**Figure 3:**
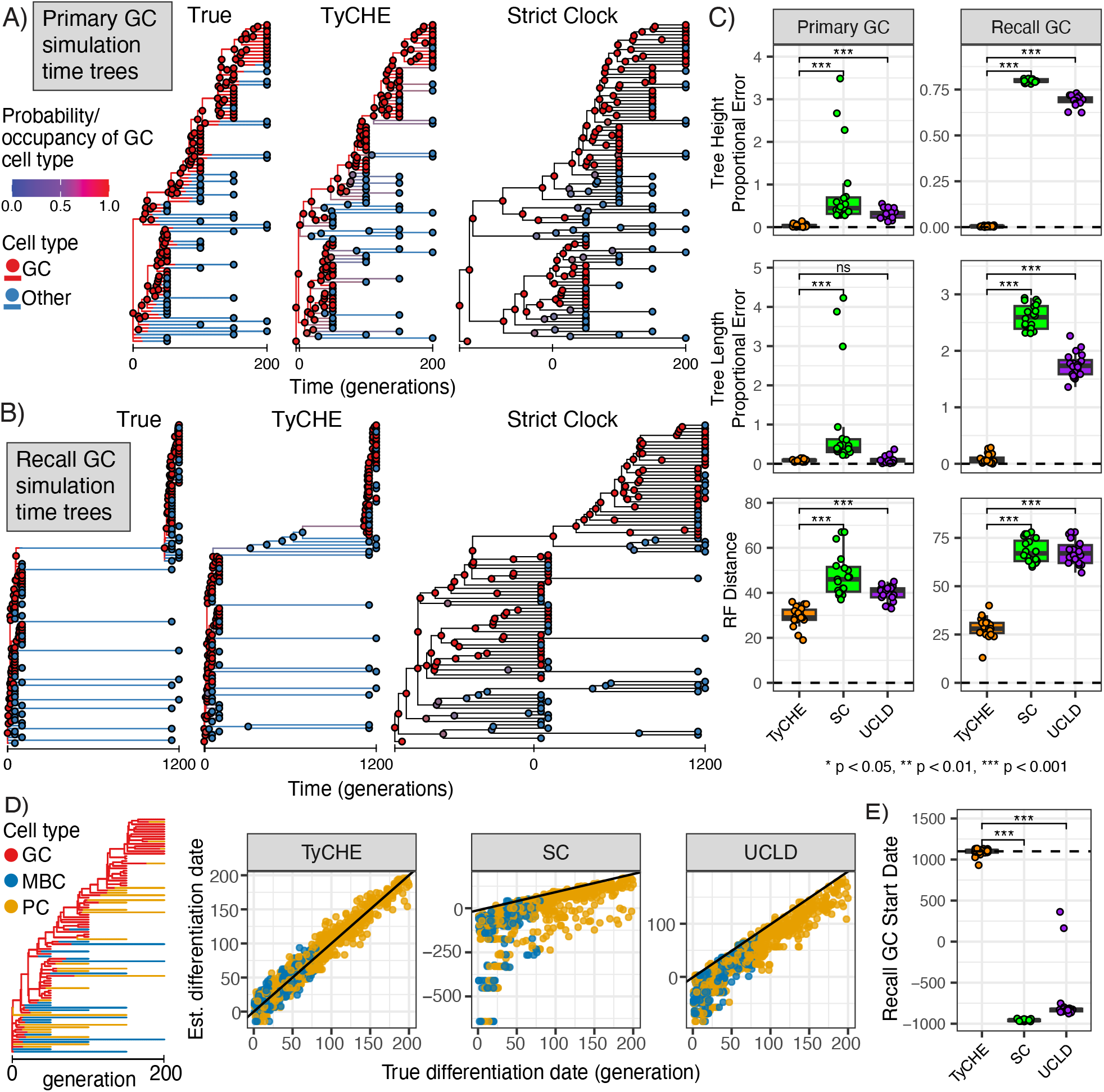
Simulation-based validation of TyCHE using SimBLE. **A)** Example results from one simulated primary GC lineage over 200 generations. From left to right: true time tree, time tree estimated by TyCHE, and time tree estimated using the SC model. All models assumed switches from GC B cells to other B cells are irreversible. **B)** Example results from one simulated recall GC lineage, in which a primary GC occurred for 100 generations before switching to memory B cells, one of which formed a recall GC at generation 1100 which was sampled for 100 more generations. From left to right: true time tree, time tree estimated by TyCHE, and time tree estimated using the SC model. All models allow switches from GC B cells to other B cells. **C)** Benchmarking of TyCHE and existing models for primary (left) and recall (right) GC simulations. Each dot represents one simulated clone. From top to bottom: proportional error of estimated tree height (generations from root to most recent tip), dashed line at 0 represents no error; proportional error of tree length estimates, calculated as the sum of all branch lengths, dashed line at 0 represents no error; Robinson-Foulds distance from the true tree topology, dashed line at 0 represents identical topology to the true tree. To reduce noise, branches under 5 generations were collapsed for Robinson-Foulds distance calculation. P values were calculated using the paired Wilcoxon signed-rank test. **D)** Using TyCHE to estimate the date of differentiation for memory (MBC) and plasma (PC) cells. Left: example true tree from SimBLE showing cell type at each generation on each branch. Right: Estimated date of differentiation from each model compared to the true date. Black line represents perfect accuracy. **E)** Estimated date of recall GC initiation during GC recall simulations (see panel B). Dashed line shows the true GC recall date at generation 1100. P values were calculated using the Wilcoxon signed-rank test.

Understanding the dynamics of cell differentiation is an important potential application of cellular time trees. We tested TyCHE’s ability to estimate MBC and plasma cell differentiation times from primary GCs. After estimating tree topologies and node heights, we traced each sampled non-GC B cell to its most recent GC ancestor. We estimated the date of differentiation using the date of this GC ancestor, adjusted by the expected occupancy in the GC state for its descendant branch. TyCHE accurately reconstructed the date of differentiation for MBCs and plasma cells, far outperforming estimates from existing models (**Fig. 3D**). Another important application is identifying when recall GCs initiated in response to a perturbation, for example a vaccine. We tested TyCHE’s ability to infer initiation dates of GC recall reactions, defined as contiguous set of internal nodes predicted to be GC B cells leading to sampled GC B cells. Across all simulations, TyCHE accurately predicted the initiation date of each recall GC reaction, approximately 1000 generations after the primary GC (**Fig. 3E**). These results confirm that TyCHE can accurately estimate the timing of cellular differentiation events out of primary GCs, and then back in to recall GCs.

### Chronic GC reactions during HIV-1 infection

HIV infection stimulates chronic GC reactions with high levels of atypical MBCs, which have been associated with autoimmune disease.^32,55–57^ To understand the dynamics of MBC differentiation during HIV infection, we investigated a dataset of bulk BCR sequences from lymph node biopsies of 3 HIV-1 infected donors clinically estimated to have been infected 4 months to 3 years prior based on history of HIV testing and antibody titers (**Fig. 4A**, donor H1: 12-18mo, H2: 24-36mo, H3: 4-8mo; Dr. Susan Moir, personal communication).^32^ B cells were sorted into GC B cells (GCBC), unswitched MBC (UnMem), CD19^lo^ (classical MBC), and CD19^hi^ (atypical MBC). Prior work has suggested that these atypical MBCs exit GC reactions prior to classical MBCs, making them potentially less able to effectively respond to infection.

**Figure 4:**
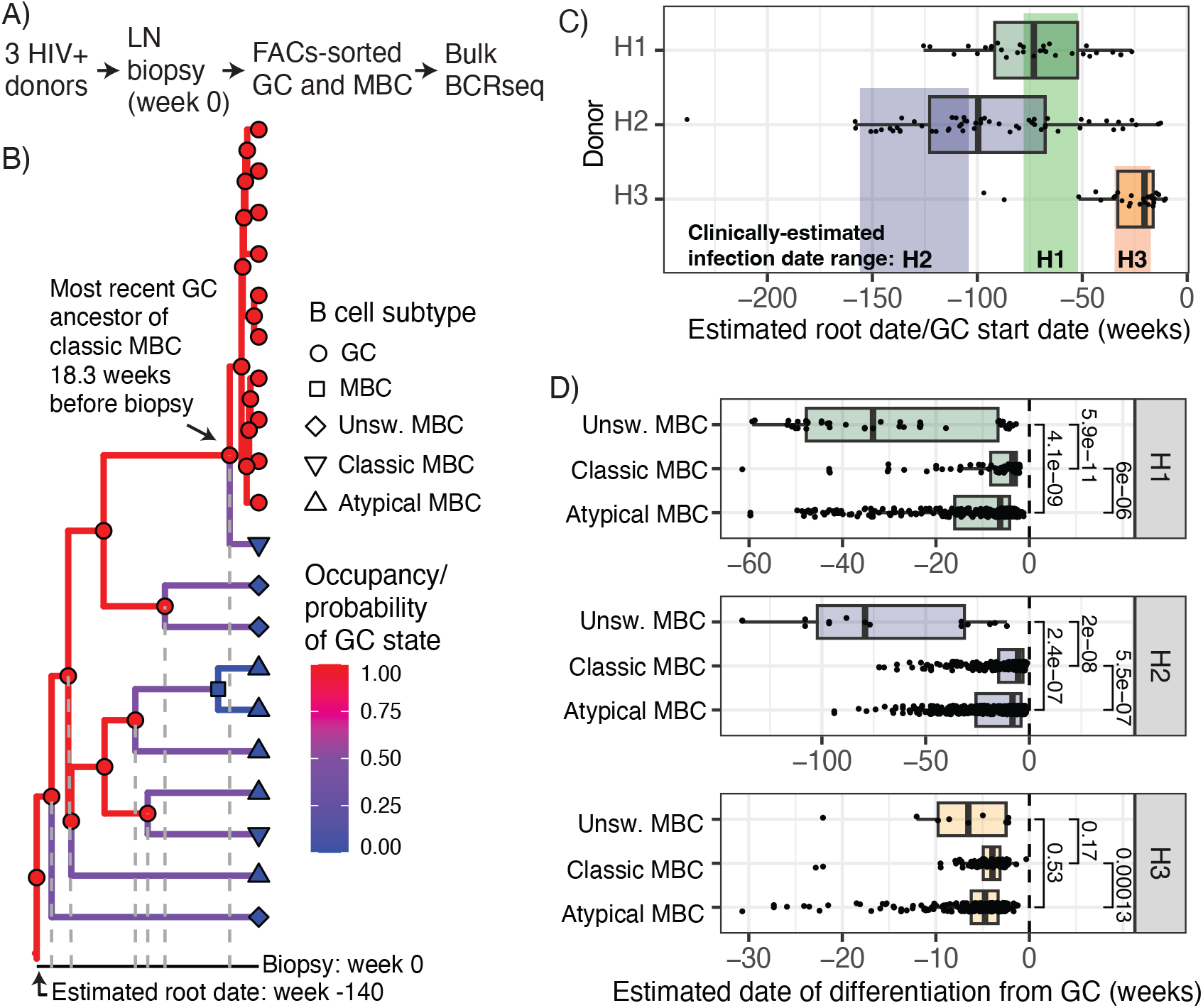
Time-resolved analysis of GC reactions in chronic HIV infection. **A)** Sampling strategy used in ^32^. **B)** Example time tree estimated by TyCHE, showing observed B cell subtypes of tips and probability of GC/non-GC type at each internal node, with branches colored by expected occupancy as GC between nodes. The estimated date of the root node, which indicates the date that this lineage began evolving in the GC, is shown. Also shown is the most recent GC ancestor of each sampled MBC (grey lines), which combined with the expected occupancy of its descendant branch indicates the estimated date of differentiation from the GC. **C)** Estimated root dates (GC initiation date) for B cell lineages sampled from each donor. Each dot represents a lineage. Clinically-estimated range of infection dates for each patient are shown in colored bands. **D)** Estimated date of differentiation from the GC (panel B) for each MBC subtype. Each dot represents a sampled MBC. The date of biopsy (week 0) is shown as a dashed line.

Each donor was sampled at a single timepoint, meaning type-specific clock rates had to be estimated externally. To obtain a GC clock rate estimate, we used previously identified measurably evolving B cell lineages from an HIV-1 infected donor sampled between 1 and 3 years of HIV-1 infection.^47,58^ We estimated the clock rate of each lineage using root-to-tip regression; i.e., the slope of the linear regression of sample time vs genetic divergence from the most recent common ancestor (MRCA).^58,59^ We used the mean rate of these 118 lineages (0.001 mut/site/week) as the mean of a strong clock rate prior for GC B cells and an arbitrarily low clock rate of 10^−6^ mut/site/week for MBCs.

We used TyCHE to reconstruct the timing and MBC differentiation patterns from sampled GC reactions among the three donors (**Fig. 4B**). As these were likely primary GC reactions, we disallowed cell type switches from non-GC B cells to GC B cells. Using TyCHE, we estimated the root date of each lineage, representing the initiation date of each GC reaction (**Fig. 4B**). Strikingly, median estimated root dates were either during or shortly after the clinically estimated time of infection for all three donors (**Fig. 4C**). This indicates that the sampled GC reactions likely began shortly after infection and persisted for months to years afterwards, consistent with other studies showing ongoing B cell evolution during chronic HIV infection.^47,55,60^ This result further demonstrates that TyCHE can use real BCR sequences to estimate biologically plausible initiation dates for primary GC reactions.

To understand the dynamics of MBC differentiation from chronic GCs during HIV infection, we estimated the date of differentiation for each sampled MBC (**Fig. 4B**, similarly to **Fig. 3D**). In all three donors, we found that unswitched MBCs differentiated from the GC earliest (significantly so in H1 and H2), followed by CD19^hi^ atypical MBCs and finally CD19^lo^ classical MBCs (**Fig. 4D**). In all three donors, atypical MBCs differentiated from the GC significantly earlier than classical MBCs. This result is consistent with prior reports that these atypical MBCs exit the GC early and accumulate in lymph nodes (**Fig. 4D**), possibly impairing viral response. These results demonstrate that TyCHE can use real BCR sequences accurately predict the relative timing of MBC differentiation patterns from primary GCs.

### Recall GC reactions following repeated human influenza vaccination

In contrast to HIV infection, which generates chronic GC reactions, influenza vaccination in adults elicits a memory response in which MBCs can form recall GCs and undergo additional evolution.^33,48,58,61^ Thus, these lineages experience periodic bursts of evolution during recall GCs preceded and followed by years of little evolution in MBCs. We used TyCHE to investigate a published dataset of single cell BCR sequences from longitudinal blood, lymph node, and bone marrow samples obtained from two human adults following seasonal influenza vaccination over the 2018/2019 and 2019/2020 flu seasons (**Fig. 5, Supplemental Fig. 5-6**).^33^ Among these data we identified 8 lineages that contained GC B cells sampled after each vaccination and were experimentally confirmed to bind to influenza antigens using monoclonal antibodies. We used a mean clock rate of 4.9 × 10^−3^ mut/site/week for GC B cells in this analysis, which was the median slope from root-to-tip regressions of previously-identified measurably evolving lineages from one of these donors (P05) sampled for 60 days after the 2018/2019 flu vaccine.^58^ To help with convergence and identifiability of the reversable model (**Supplemental Fig. 3**), the cell type at the root was fixed to GC and the maximum root date of the trees was set to four years prior to the 2018 QIV vaccination for all models, as neither patient had received a flu vaccine in the preceding three years. This dataset provides a unique opportunity to understand the timing and differentiation dynamics of recall GCs in humans.

**Figure 5:**
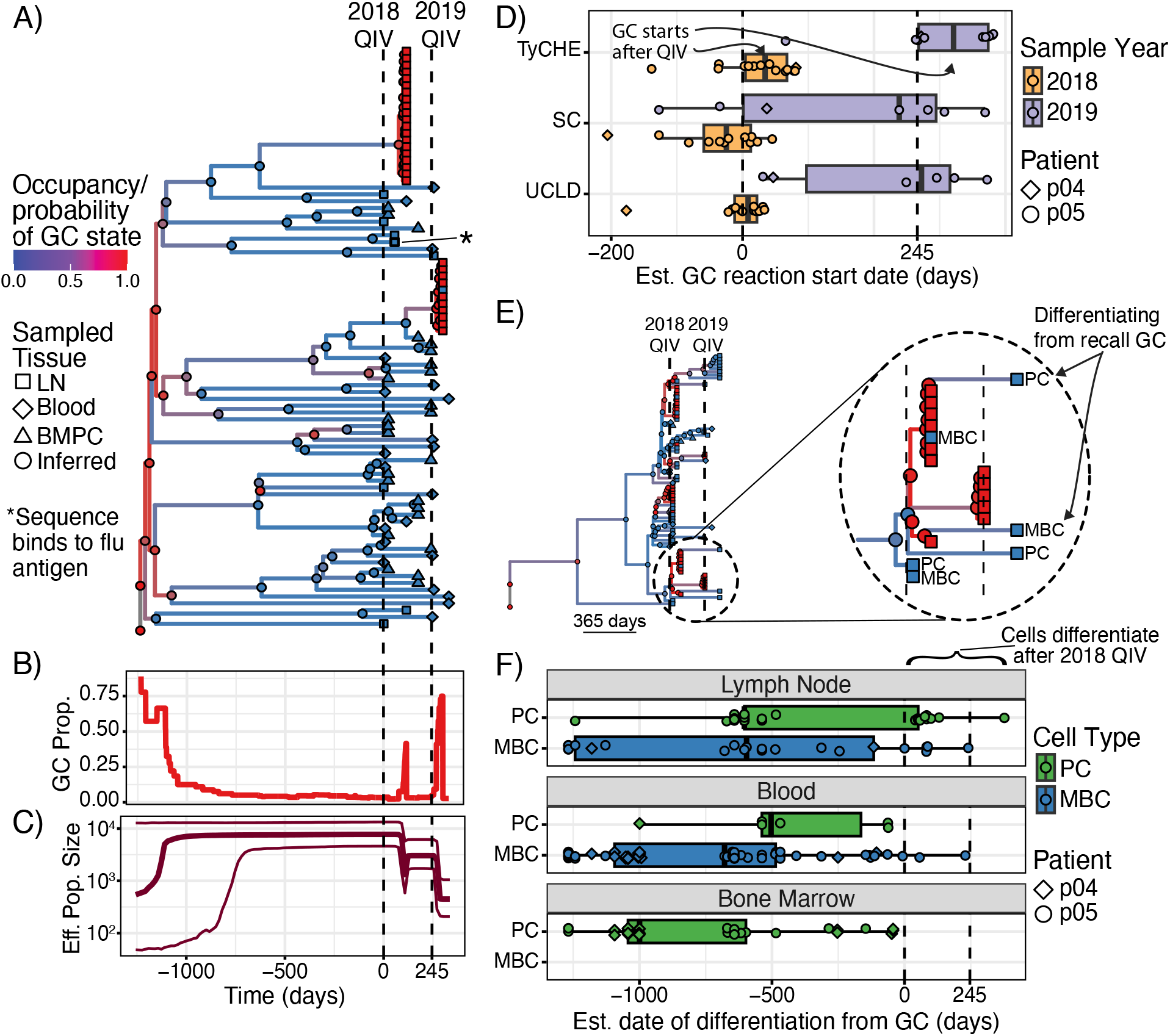
Time-resolved analysis of recall GC reactions following repeated influenza vaccination. Donors were immunized with 2018/2019 QIV at week 0 and re-immunized with 2019/2020 QIV at week 35 (P04) or 38 (P05). Samples were taken from peripheral blood, lymph node (LN), and bone marrow plasma cells (BMPCs). **A)** TyCHE estimated time tree of the largest lineage from P04. Dashed lines show dates of vaccinations. **B)** Proportion of branches in the above tree predicted to be GC B cells. **C)** Bayesian skyline plot showing estimated changes in effective population size for the clone in panel A. **D)** Estimated recall GC reaction start dates. Each dot represents a sampled GC reaction. Boxplots and dots are colored by the flu season in which GC B cells were sampled. Dotted lines show the dates of 2018 and 2019 QIVs for P05. **E)** Example time tree showing MBC and PC in the LN produced by a sampled recall GC. Legend as in panel A. **F)** Estimated date of differentiation for plasma (PCs) and memory (MBCs) cells by sample tissue. Dots represent sampled B cells. Dotted lines show the dates of 2018 and 2019 QIVs for P05. Dots shown between these lines represent MBCs and PCs produced by recall GC reactions from the 2018 QIV.

We tested whether TyCHE could infer the periodicity of recall GC reactions using these data. TyCHE correctly predicted bursts of GC B cells following vaccination with long periods of MBCs in between (**Fig. 5A-C, Supplemental Fig. 5-6**). We defined a recall GC reaction as a contiguous set of internal nodes predicted to be GC B cells leading to sampled GC B cells, and its initiation date as the date of its earliest GC internal node. These recall GCs were presumably induced by the vaccine and should have begun only after vaccination. TyCHE correctly inferred that these recall GCs began at or after the date of each season’s vaccination 79% of the time (**Fig. 5D**). By contrast, SC and UCLD (**Supplemental Fig. 7**) models predicted that only 42% and 50% (respectively) of these GC reactions began after each season’s vaccination, which is biologically implausible (**Fig. 5D**). Even less plausibly, the SC model predicted that 29% of GCs sampled after the 2019/2020 vaccination began before even the 2018/2019 vaccination (**Fig. 5D**, purple boxplots). Recall GCs were also detectable using changes in effective population size. Using a Bayesian skyline analysis of the largest clone, TyCHE inferred sharp decreases in effective population size during recall GCs, consistent with strong selection and clonal expansion (**Fig. 5C**). These results demonstrate that TyCHE can use real BCRs to accurately reconstruct the timing and dynamics of recall GCs.

Understanding when and how GCs generate MBC and plasma cells, particularly long-lived plasma cells in the bone marrow, is critical for designing better vaccines. We used TyCHE to estimate the differentiation dates for each of these cell populations in each tissue. First, the time tree of one clone showed clear evidence of recall GCs in the 2018/2019 season producing MBCs and plasma cells that persisted in the LN beyond the 2019/2020 vaccine (**Fig. 5E**). This confirms that recall GCs can produce these effector populations. However, these cells were rare, and no plasma cells originating from sampled recall GCs were found in the blood or bone marrow (**Fig. 5F**). B cells sampled in the blood followed a classic pattern of MBCs differentiating earlier than plasma cells. However, bone marrow plasma cells were estimated to differentiate earlier on average than all other cell types studied, indicating they are more likely to be produced by earlier, perhaps primary, GC reactions (**Fig. 5F**). If so, this could be a straightforward mechanism of observations of “original antigenic sin” – that recall GCs have reduced capacity to produce long-lived bone marrow plasma cells.

### Evolution of a hypermutating glioma tumor lineage

Tumors frequently accumulate mutations over time, and phylogenetic analysis of these mutations can reveal significant insights into the response of tumors to treatment.^8^ Some tumors develop hypermutator phenotypes, which frequently occur in gliomas during temozolomide treatment. We used TyCHE to resolve the evolution of a mixed hypermutator (H)/non-hypermutator (N) glioma lineage in one patient profiled in the GLASS consortium (**Fig. 6A**).^34^ Samples included normal genome (NG), primary tumor (TP, day 0), recurrence 1 (R1, day 3042) and recurrence 2 (R2, day 3772). R2 was classified as a hypermutator based on mutation levels (**Fig. 6B**). We estimated the clock rate of N cells using the SC model on NG, TP, and R1, as well as the clock rate of H using root-to-tip regression of R1, R2, and NG. Using the SC model on only N sequences, we estimated the root date of the lineage at ∼15.1 years before TP, consistent with slow-growing low-grade gliomas (**Fig. 6C**). Using all sequences, TyCHE estimated a similar root date of 16.5 years prior to TP (**Fig. 6C-D**). By contrast, the SC on all sequences estimated an implausibly recent root date of 53.5 days prior to TP, likely due to an inflated clock rate estimate, while the UCLD model predicted an implausibly early root date of 58.9 years before TP, >20 years before the patient was born (**Fig. 6C**). These results demonstrate that TyCHE can reconstruct the temporal dynamics of complex, heterogeneous tumor populations with hypermutating subclonal populations.

**Figure 6:**
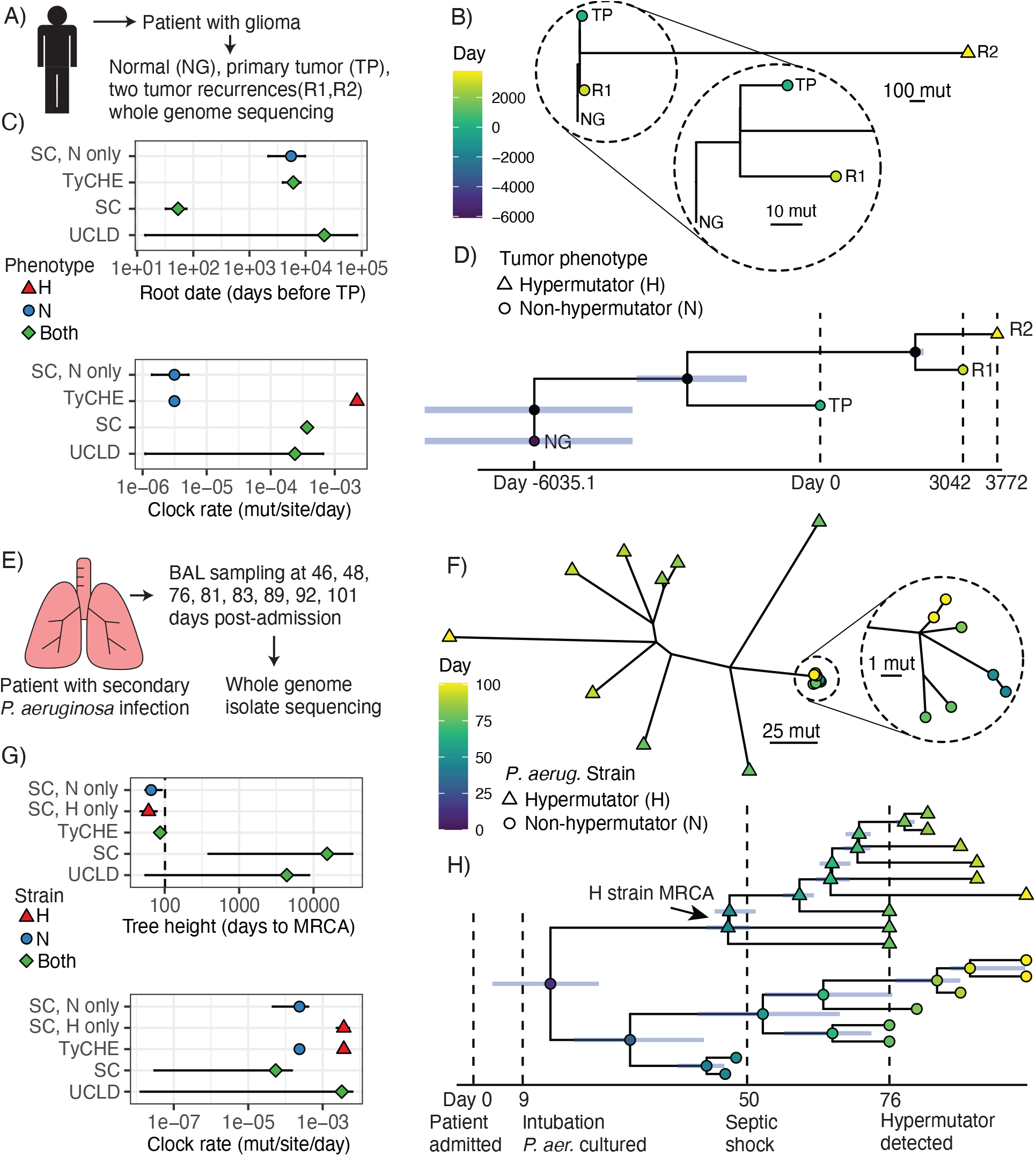
TyCHE analysis of a hypermutating glioma tumor lineage (A-D) and a *Pseudomonas aeruginosa* lung infection (E-H). **A)** Tumor genome sampling from a patient with glioma. **B)** Genetic distance tree of hypermutator (R2) and non-hypermutator (TP, R1) tumors rooted at NG. Non-hypermutator branches zoomed in for detail. This panel shares color/shape scales with panel D. **C)** Root date (above) and clock rate (below) estimates from each model, both in log_10_ scale. **D)** Time-resolved tumor phylogeny from TyCHE, tip color and shape shares scales with panel B. Blue bars show 95% highest posterior density (HPD) interval of node height. **E)** Bronchial alveolar lavage (BAL) sampling from a patient with COVID-19 associated pneumonia and secondary *P. aeruginosa* infection. **F)** Unrooted genetic distance tree of hypermutator (H) and non-hypermutator (N) isolates. N clade zoomed in for detail. This panel shares color/shape scales with panel H. **G)** Tree height (above) and clock rate (below) estimates from each model, both in log_10_ scale. Tree height of ∼101 days (dashed line) is expected as the patient showed no signs of lung infection before hospitalization. **H)** Time-resolved phylogeny of H and N isolates, shares color/shape scales with panel F. Blue bars show 95% HPD interval of node height. MRCA of all isolates estimated to be at/shortly after intubation. MRCA of H estimated shortly before the septic shock, and H lineage evolved in a ladder-like manner shortly afterwards.

### Evolution of a hypermutating bacterial lung infection

*P. aeruginosa* can cause deadly lung infections which often begin with a single bacterium that rapidly adapts to the lung environment.^24^ Mutations in DNA repair enzymes produce hypermutator lineages that evolve far more rapidly than wild-type strains.^25,62^ We used TyCHE to reconstruct the evolution of a *P. aeruginosa* hypermutator lineage in an acute, fatal lung infection.^25^ Whole genome sequencing was previously performed on isolates obtained from bronchial alveolar lavage samples of a patient admitted for COVID-19 related pneumonia with no indication of prior *Pseudomonas* infection (**Fig. 6E**). *P. aeruginosa* was first detected by culture on day 9, and on day 76 a hypermutator (H) strain was detected with genetic diversity far exceeding that of non-hypermutator (N) isolates (**Fig. 6F**). Using the SC model on H and N strains separately, we estimated similar MRCA dates of 40.3 and 35.4 days post-admission, respectively, but a much higher clock rate for H (**Fig. 6G**). Using these clock rate estimates as prior means for H and N strains, respectively, TyCHE estimated an MRCA date of all sequences at 14.1 days post-admission with a 95% HPD (3.1 - 24.2 days) overlapping with the patient’s first positive *P. aeruginosa* sputum sample at intubation on day 9 (**Fig. 6H**). By contrast, SC and UCLD clock models estimated implausibly early root dates of 41.4 years and 11.7 years before admission, respectively (**Fig. 6G**). Using TyCHE, we further estimated the MRCA of the H strain at day 46.5, 4 days before the patient experienced septic shock on day 50. The H strain also exhibited a ladder-like branching pattern, consistent with rapid evolution, roughly corresponding to a two-week period of intense antibiotic treatment following septic shock (**Fig. 6H**).^25^ These results are consistent with the H strain contributing to septic shock in this patient. Supporting this conclusion, peripheral blood samples taken at days 76, 78, and 79 contained only H strain isolates.^25^ These results confirm that TyCHE can infer realistic and clinically useful time-resolved phylogenies in diverse, hypermutating bacterial populations.

## Discussion

Inferring time-resolved phylogenies of cellular lineages could uncover new discoveries about how cell proliferation, differentiation, and migration shape responses to perturbations like vaccines and drug treatments. Unfortunately, mutation rates often vary significantly by cell type, biasing existing models. Here, we introduce TyCHE, which uses type-linked clock models to estimate accurate time trees for heterogeneously evolving populations. Using simulations, we show TyCHE estimates more accurate tree topologies, node dates, and cell differentiation times than existing methods. Further, we show how TyCHE can use BCR sequences to reconstruct both primary GC reactions in HIV infection and recall GC reactions following influenza vaccination. When applied to sequences from hypermutating tumor lineages and bacteria, TyCHE can infer biologically plausible and clinically useful time trees. By contrast, existing SC and UCLD models produced biologically implausible results. Further, we introduce SimBLE, which simulates both BCRs evolving during GC reactions and neutral heterogeneous evolution, to benchmark these new models and perform *in silico* evolution experiments.

While most phylogenetic models that allow clock rate heterogeneity allow rates to vary independently of cell type,^30,36^ some prior studies have linked evolutionary rate with discrete traits. One study introduced a likelihood-based method to detect shifts in trait-dependent evolution.^63^ More recently, the package SDevo linked tumor cell birth rates in a birth-death model to discrete states, specifically the position of cancer cells along the edges of a tumor.^64^ TyCHE differs from SDevo’s approach by allowing clock rates rather than birth rates to vary by cell type, introducing the expected occupancy approximation, and by utilizing a simpler CTMC model. One limitation of TyCHE (though not unique to TyCHE) is that prior information about clock rates is usually needed, but this can be obtained using strict clock models or root-to-tip regressions applied to pure cell populations, as was done in this study. In cases where only one timepoint is sampled, these rates can be estimated externally, as done in **Fig. 4**. Because of this two-step process, similar to empirical Bayes methods, 95% highest posterior density intervals should always be interpreted as conditioned on the provided cell-type clock rates, especially if estimated from the same data. Further, adequate and biologically plausible model constraints are often needed to find identifiable posterior estimates. Assuming irreversibility of type switches is highly effective when biologically justified (e.g. **Figs. 3, 4**, and **6**). While this is not plausible in recall GCs, we show that fixing the root cell type and setting a plausible maximum tree height work well in practice (**Figs. 3** and **5**). The examples tested in this study used cell types with rates differing by orders of magnitude. SC and UCLD models may be more competitive when analyzing cell types with more similar rates. While in this study we focused on naturally-occurring somatic mutations, TyCHE could be applied to dynamically mutating lineage recorders if combined with an appropriate sequence substitution model (e.g. ^21,22,65^).

We introduce TyCHE and SimBLE for accurately reconstructing and simulating heterogeneously evolving populations, respectively. Because type-linked mutational heterogeneity is a very general phenomenon, we believe the examples explored in this study are a small fraction of potential applications for these programs.

## Accessibility

TyCHE is available as a BEAST2 package on GitHub: https://github.com/hoehnlab/tyche. SimBLE is also available on GitHub and PyPi: https://github.com/hoehnlab/simble. Both interface with the R package Dowser for ease of use: https://dowser.readthedocs.io. Scripts and BEAST2 XML templates to reproduce these analyses are available on GitHub: https://github.com/hoehnlab/publication_scripts. All empirical data are publicly available from prior publications.

## Acknowledgements

This work was supported by NIH grants R00AI159302 and R35GM160119 (K.B.H). H.J.M. was supported by NIH grant T32CA134286. We thank Drs. Alan Hauser and Egon Ozer for providing *P. aeruginosa* sequence alignments, Dr. Susan Moir for providing estimated dates of infection of HIV+ donors, Dr. Aaron McKenna for helpful comments, and Cole Jensen for early testing of SimBLE.

## Methods

TyCHE (Type-linked Clocks for Heterogenous Evolution) introduces a framework for time-resolved Bayesian phylogenetic inference on populations that evolve at different rates. TyCHE is implemented as a new BEAST2^31^ package and validated with SimBLE (Simulator of B cell Lineage Evolution). As with other BEAST2 packages, the type-linked models in TyCHE are specified using XML files. For ease of use, especially on multiple lineages in concert, we developed a system of functions within the R package Dowser^66^ that allows users to specify general XML templates applied to multiple lineages at once. A vignette showing how TyCHE can be used on both B cells and non-B cells is available at https://dowser.readthedocs.io.

### Type-linked clock model

The observed data comprise discrete states (e.g. cell types) ***X*** = (*X*_1_, …, *X*_*N*_) and aligned nucleotide sequences ***Y*** = (*Y*_1_, …, *Y*_*N*_) at the *N* tips of the phylogenetic tree ***F***. For each tip, we observe a state *X*_*i*_ ∈ *S*_*X*_ and a sequence *Y*_*i*_ ∈ *S*_*Y*_, where *S*_*x*_ is the set of all states and *S*_*y*_ is the set of all potential nucleotide sequences of common length *L*. Neither states nor sequences are observed at the internal nodes of the phylogeny or the root; i.e. *X*_*root*_, *Y*_*root*_, (*X*_*N*+1_, …, *X*_2*N*−2_), (*Y*_*N*+1_, …, *Y*_2*N*−2_) are unknown.

We adapt a Bayesian discrete ancestral state reconstruction method^39^ to model the state changes *X*(*t*) along the branches of the phylogeny ***F*** via continuous time Markov chains (CTMCs). CTMCs are stochastic processes in continuous time that emit discrete outcomes and are memoryless; i.e., ∀*i, j* ∈ *S*, ∀*s, t* ≥ 0, P(*X*(*s* + *t*) = *j* | *X*(*s*) = *i*, {*X*(*u*): 0 ≤ *u* ≤ *s*}) = P(*X*(*s* + *t*) = *j* |*X*(*S*) = *i*). A two-state CTMC is fully specified by the instantaneous rate matri 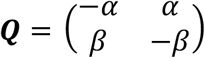, which necessarily contains non-negative off-diagonal entries and rows that sum to 0. Solving the Chapman-Kolmogorov equation results in the finite-time state transition matrix **P**(*t*) = *e*^***Q****t*^ = {P_*ij*_(*t*)}, where P_*ij*_(*t*) = P(*X*(*t*) = *j* | *X*(0) = *i*). The sequences ***Y*** are similarly modeled with a CTMC which assumes each site mutates independently according to the instantaneous rate matrix ***Q***_*Y*_, resulting in the joint TyCHE Bayesian hierarchical model (BHM):

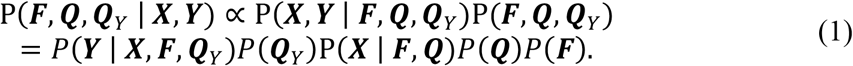

The phylogeny prior, *P*(***F***), can be chosen to be any valid tree prior. In this work, we use a Bayesian skyline tree prior^67^ in all analyses and validations unless otherwise specified.

The state instantaneous rate matrix ***Q*** can be decomposed into the product of the type-switch clock rate *μ*_*X*_, and a normalized instantaneous rate matrix 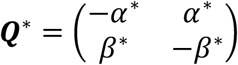. The hyperparameter *α*^∗^ = *r*_*AB*_*f*_*A*_, where *r*_*AB*_ is the relative rate of transition from state *A* to state *B* and *f*_*A*_ is the relative frequency of state *A*. The hyperparameter *β*^∗^ = *r*_*BA*_*f*_*B*_ is defined analogously. Therefore, the prior distribution of the state instantaneous rate matrix ***Q*** can be written *P*(***Q***) = *P*(*μ*_*X*_)*P*(*r*_*AB*_)*P*(*f*_*A*_)*P*(*r*_*BA*_)*P*(*f*_*B*_). The relative frequencies of the states in our analyses are likely changing over time. As such, we wish to minimize the impact of the relative frequencies in the model, and set *f*_*A*_ = *f*_*B*_ = 0.5 to avoid biasing *Q*^∗^.

The typed-tree likelihood, P(***X*** | ***F, Q***), represents the likelihood of the internal node states given the observed states at the tips, the phylogeny ***F***, and the state instantaneous rate matrix ***Q*** = *μ*_*X*_ ∗ ***Q***^∗^. It is assumed that the sequence CTMC depends on the internal node states, but the state CTMC is conditionally independent of the sequences given the phylogeny ***F***. The typed-tree likelihood can be decomposed into the product of the state transition probabilities along each branch *i* of ***F***, P(***X*** | ***F, Q***) = Π_*i*_ *P*(*X*_*child*_|*X*_*parent*_, *t*_*i*_, ***Q***) = Π_*i*_ *P*(*X*_*child*_|*X*_*parent*_, *t*_*i*_ ∗ *μ*_*X*_, ***Q***^∗^), where *X*_*child*_ is the state of the child node on branch *i, X*_*parent*_ is the state of the parent node on branch *i*, and *t*_*i*_ is the length in time of branch *i*; *μ*_*X*_ is the type-switch clock rate such that the product *t*_*i*_ ∗ *μ*_*X*_ represents the distance between parent and child types, and ***Q***^∗^is the normalized instantaneous rate matrix. This represents the joint likelihood of state transitions given the states at each internal node, and differs from other approaches^39^ which assume node states do not affect clock rates and therefore sample states from the marginal posterior probabilities at each node.

The nucleotide substitution instantaneous rate matrix ***Q***_*Y*_ is specified by the chosen nucleotide substitution model. Unless otherwise noted, the HKY substitution model^68^ was used with equilibrium nucleotide frequencies fixed to their empirical frequencies, and a prior was placed on the transition/transversion rate ratio *κ*, so *P*(***Q***_*Y*_) = *P*(*κ*).

The tree likelihood *P*(***Y*** | ***X, F, Q***_*Y*_) is calculated as a standard BEAST2 tree likelihood.^31^ Briefly, a clock model is used to convert the branches of the tree ***F*** into a genetic distance tree, and the likelihood of that tree is calculated using Felsenstein’s pruning algorithm.^37^ As in a relaxed clock model^30^ or random local clock model,^36^ in TyCHE each branch receives an individual rate of evolution, or branch rate, which is defined as the rate of genetic mutation for the sequences ***Y*** along a given branch of the phylogeny ***F***. Unlike in previous models, in TyCHE the branch rate is linked to the state of the parent and child nodes.

For ease of notation, let the binary state space *S*_*X*_ = {*A, B*}, and denote the state-specific mutation rates as *μ*_*A*_ and *μ*_*B*_. We present TyCHE’s type-linked evolutionary rate expected occupancy time (EO) model below.

The EO model is implemented for a two-state case and determines the expected amount of time spent in each state along a branch as a function of the parent and child node states and the instantaneous rate matrix ***Q***. Then the overall branch rate is

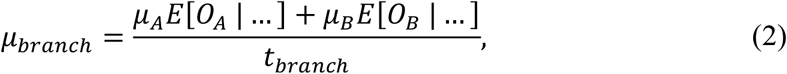

where *E*[*O*_*A*_ | …] is the expected occupancy time in state *A* given the parent state, child state, and instantaneous rate matrix ***Q***, *E*[*O*_*B*_ | …] is analogous for state *B*, and *t*_*branch*_ is the length of the branch in time. Note that *t* = *E*[*O*_*A*_ | …] + *E*[*O*_*B*_ | …], so this is a weighted average of the state-specific mutation rates. The derivation of the expected occupancy times is given in the following section. We provide the option for users to either estimate the state-specific mutation rates or provide fixed values.

### Calculating the expected occupancy time

In TyCHE’s expected occupancy time model, the expected occupancy time in each state along each branch of the phylogeny ***F*** is determined by the instantaneous rate matrix 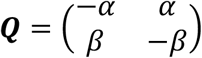, the time length of the branch, and the parent and child states. For ease of notation, let the binary state space *S*_*X*_ = {*A, B*}, and set the time such that *t*_*parent*_ = 0 and *t*_*child*_ = *t*. Then the state transition matrix is along the branch is 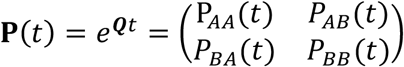, where P_*ij*_(*t*) = P(*X*(*t*) = *j* | *X*(0) = *i*) ∀*i, j* ∈ *S*_*x*_.

As an example, let *X*(0) = *A, X*(*t*) = *B*, i.e., the parent state is *A* and the child state is *B*. Per ^69^ the occupancy time in state *A* can be written as

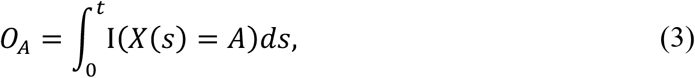

where I(·) is the indicator function. Then the expected value of the occupancy time in state A given the parent and child states is

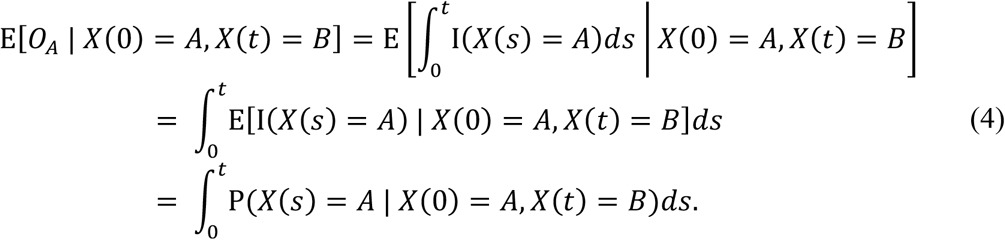

In these expressions, the order of the expectation and integration are interchanged by Fubini’s theorem, and the expectation of the indicator function is the simple probability. Consider now the expression P(*X*(*s*) = *A* | *X*(0) = *A, X*(*t*) = *B*). By a conditioning argument,

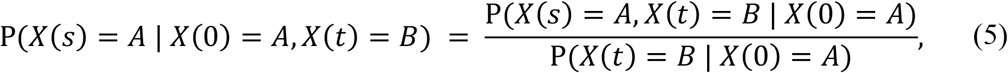

and P(*X*(*t*) = *B* | *X*(0) = *A*) = P_*AB*_(*t*). By a similar conditioning argument,

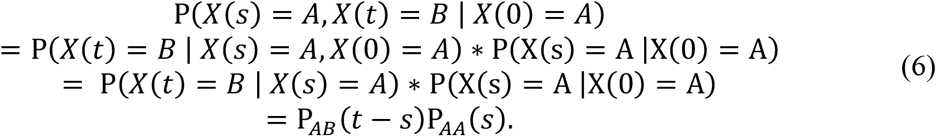

Therefore,

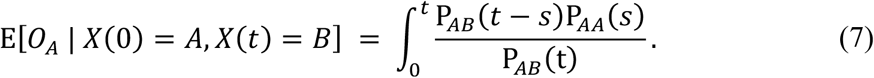

To solve this expression, it is necessary to determine the forms of each of the functions in the state transition matrix 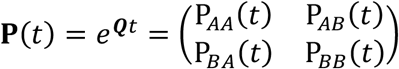. Consider the eigen-decomposition ***Q*** = ***VDV***^−1^. The eigenvalues of ***Q*** are *λ*_1_ = 0, *λ*_2_ = −(α + β), and the eigenvectors of ***Q*** are taken to be 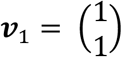 and 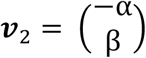, so 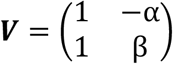 and 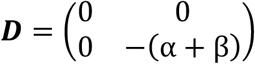. Then,

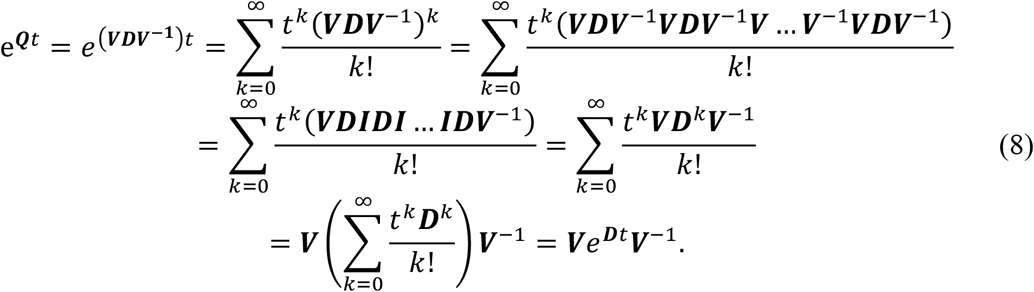

As ***D*** is diagonal,

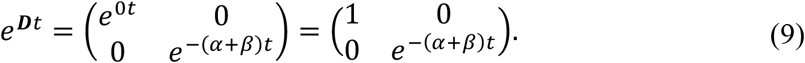

Thus,

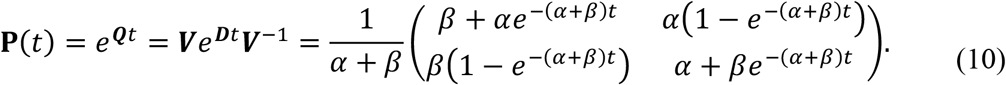

Inserting these expressions into Equation (7), the expected occupancy time is

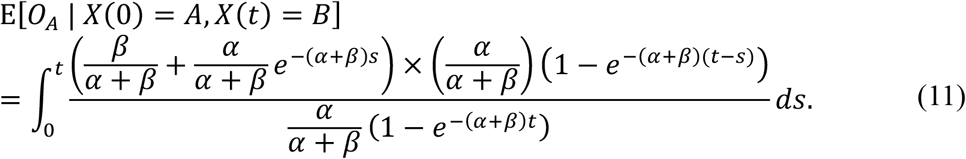

Solving the integral analytically leads to the result

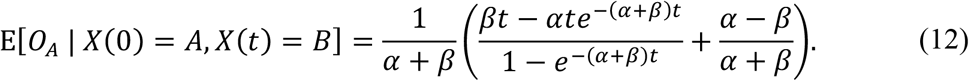

While the occupancy time in state *B* can be solved for similarly, it is simpler to consider that, for a two state sample space *S*_*X*_,

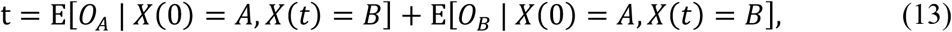

and thus,

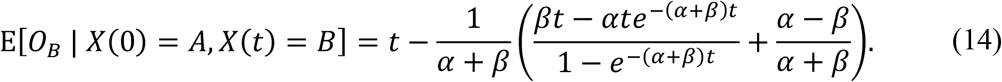

Expressions for different combinations of parent states *X*(0) and child states *X*(*t*) can similarly be derived and are presented in **Table 1**.

**Table 1:** Expected occupancy times given starting and ending states along a branch of length *t*.

| $X(0)$ | $X(t)$ | Expected Occupancy Time |
| --- | --- | --- |
| A | B | $E[O_A \vert X(0) = A, X(t) = B] = \frac{1}{\alpha + \beta} \left( \frac{\beta t - \alpha t e^{-(\alpha+\beta)t}}{1 - e^{-(\alpha+\beta)t}} + \frac{\alpha - \beta}{\alpha + \beta} \right)$ |
| | | $E[O_B \vert X(0) = A, X(t) = B] = t - \frac{1}{\alpha + \beta} \left( \frac{\beta t - \alpha t e^{-(\alpha+\beta)t}}{1 - e^{-(\alpha+\beta)t}} + \frac{\alpha - \beta}{\alpha + \beta} \right)$ |
| B | A | $E[O_A \vert X(0) = B, X(t) = A] = \frac{1}{\alpha + \beta} \left( \frac{\beta t - \alpha t e^{-(\alpha+\beta)t}}{1 - e^{-(\alpha+\beta)t}} + \frac{\alpha - \beta}{\alpha + \beta} \right)$ |
| | | $E[O_B \vert X(0) = B, X(t) = A] = t - \frac{1}{\alpha + \beta} \left( \frac{\beta t - \alpha t e^{-(\alpha+\beta)t}}{1 - e^{-(\alpha+\beta)t}} + \frac{\alpha - \beta}{\alpha + \beta} \right)$ |
| A | A | $E[O_A \vert X(0) = A, X(t) = A] = \frac{1}{\alpha + \beta} \left( \frac{\beta^2 t + \frac{2\alpha\beta}{\alpha + \beta} (1 - e^{-(\alpha+\beta)t}) + \alpha^2 t e^{-(\alpha+\beta)t}}{\beta + \alpha e^{-(\alpha+\beta)t}} \right)$ |
| | | $E[O_B \vert X(0) = A, X(t) = A] = t - \frac{1}{\alpha + \beta} \left( \frac{\beta^2 t + \frac{2\alpha\beta}{\alpha + \beta} (1 - e^{-(\alpha+\beta)t}) + \alpha^2 t e^{-(\alpha+\beta)t}}{\beta + \alpha e^{-(\alpha+\beta)t}} \right)$ |
| B | B | $E[O_A \vert X(0) = B, X(t) = B] = \frac{1}{\alpha + \beta} \left( \frac{\alpha\beta t - \frac{2\alpha\beta}{\alpha + \beta} (1 - e^{-(\alpha+\beta)t}) + \alpha\beta t e^{-(\alpha+\beta)t}}{\alpha + \beta e^{-(\alpha+\beta)t}} \right)$ |
| | | $E[O_B \vert X(0) = B, X(t) = B] = t - \frac{1}{\alpha + \beta} \left( \frac{\alpha\beta t - \frac{2\alpha\beta}{\alpha + \beta} (1 - e^{-(\alpha+\beta)t}) + \alpha\beta t e^{-(\alpha+\beta)t}}{\alpha + \beta e^{-(\alpha+\beta)t}} \right)$ |

### Custom priors and operators for germline-rooted time trees

To handle the germline’s type appropriately, we introduced the Germline-Root Tree class within BEAST2. This allows the germline’s relationship with the root to be fixed, such that the germline is always a child of the root, its type and sequence always match that of the root, and its node height is moved with the root height with an arbitrarily small branch length between them to avoid zero-length branches. The structure of the Germline-Root Tree requires specialized operators to ensure that this structure is never violated by tree height or topology proposals. Further, because the Germline is not a sampled sequence, the Germline-Root Tree uses a modified Bayesian Skyline tree prior that ignores the false coalescent interval between the MRCA and the root of the tree. To adequately constrain the model, for example during analysis of recall GC reactions, we introduce a root type prior, allowing the user to specify the probability of the root being a particular cell type.

TyCHE introduces several tree type operators to make bigger jumps through the state space. In particular, we introduce operators that change the type of several nodes at once and also that change the height of nodes at the same time as changing their type. This is necessary to be able to move out of local extrema resulting from type-linked models. Some of these operators cannot have a symmetrical proposal distribution, and return a Hastings ratio to ensure detailed balance.

First, a combined height and type operator for the root makes proposals by drawing a proposal type uniformly at random from the set of all types, and drawing a perturbation in height, *δ*, uniformly from a window defined as 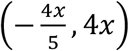, where *x* is the branch length from the root to its tallest normal child (i.e. not the germline when the tree is a Germline-Root Tree). This allows the operator to propose jumps that move the root very close or very far, which is sometimes necessary to change the type while avoiding local extrema. However, the branch length defining this window, *x*, changes with the proposal, so the operator returns the ratio of these windows as the Hastings ratio: 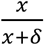. The proposal of types is symmetrical and therefore has no bearing on the Hastings ratio.

A similar combined height and type operator for two adjacent internal nodes makes proposals by picking a node *A* uniformly at random from the set of nodes that are not tips and whose grandparents are true internal nodes (i.e. not the root), and then proposing a new height for *A* uniformly between *A*’s grandparent’s height, *g*, and the height, *c*, of *A*’s tallest child. It then proposes a new height for *A*’s parent, *P*, uniformly between g and the taller of *P*’s other child, *S*, and A’s new height. Then a random type is chosen for each of *A, P*, and *S*. The proposal of types is symmetrical and therefore has no bearing on the Hastings ratio. The window in which *A* can move does not change between the starting state and the proposal state, and is therefore symmetrical and has no bearing on the Hastings ratio. The window in which *P* can move does change between the starting state and proposal state, so we return the ratio of those windows as the Hastings ratio:

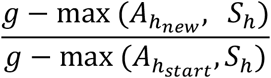

A subtree type-switch operator works by proposing types for all internal nodes and ambiguously-typed tips in the subtree rooted at a randomly chosen internal node. This operator is designed to propose large jumps to homogenous subtrees as moving a neighborhood from one homogenous type to another in individual steps often involves passing through local extrema where one node of a different type causes many more type switching events in the tree than is likely. To ensure that this operator can propose the starting state from its proposal state, it proposes homogenous subtrees half of the time and randomly heterogenous subtrees half of the time. To account for this asymmetry when the starting subtree is not entirely homogenous, we return a Hastings ratio of:

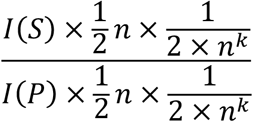

where *S* is the starting state, *P* is the proposed state, *I* represents the indicator function returning 1 when a state’s subtree is homogenous and 0 when it is not, *n* is the total number of types nodes can be, and *k* is the number of nodes in the subtree whose type can be operated on (internal nodes and ambiguous tips).

### SimBLE: realistic germinal center reaction simulations

SimBLE is an open-source Python package that simulates type-linked evolution, either neutral evolution of random sequences, or BCR sequences from B cells in GC reactions using a realistic model of SHM, affinity maturation, and differentiation. SimBLE models essential processes including B cell competition for antigen in GCs and their differentiation into MBCs and plasma cells. Briefly, Simble picks a naïve cell with paired heavy and light chain and adds it to the *in silico* GC. If simulated under selection, at each generation the cells in the GC are evaluated for relative affinity and reproduce based on that affinity. Cells migrate out of GCs as memory B cells or plasma cells at a user-specified rate, where typically they do not mutate and reproduce slowly with no selection.

To create a SimBLE naïve cell, a paired heavy and light chain are randomly selected from a previously curated dataset of naïve B cells.^35,40^ These cells were obtained from healthy donors and COVID-19 patients. Only cells transcriptionally identified as naïve B cells and containing unmutated heavy and light chain BCRs were included. The selected cell is added to the GC, where its relative fitness (affinity) is assessed based on similarity to a pair of target heavy and light chain B cell receptor amino acid sequences, similar to the approach used in *bcr-phylo*^42^ for heavy chains. The target sequence is generated by mutating the complementarity determining regions (CDRs) of the germline sequence; by default, the target sequence has 5 single nucleotide amino acid (AA) substitutions in the heavy chain and 3 in the light chain. The number of target sites may be specified by the user. Single nucleotide AA substitutions are accessible by a single nucleotide substitution from the germline state. The positions of these substitutions in the target sequence (target mutations) are drawn randomly from within the CDRs which determine antigen binding. Target mutations were not allowed in the framework regions (FWRs) because those regions are usually conserved for antibody structure.

To replicate the targeting biases of SHM, mutations are added to BCRs according to the S5F model of SHM targeting and nucleotide substitution.^41^ When a child cell is created, *n* mutations are drawn for the heavy chain from a Poisson distribution with rate = (cell type’s relative mutation rate) * (expected heavy chain mutations per site) * (number of heavy chain sites). The same process determines mutations for the light chain. By default, GC B cells have a relative mutation rate of 1, and non-GC B cells have a relative mutation rate of 0, i.e., they do not mutate. Affinity of a cell’s BCR sequence is calculated based on mismatches from the target AA sequence, with different selection multipliers (weights) for each site. A site multiplier at the lower bound of 1 represents neutral impact of that site on binding ability. A multiplier greater than 1 represents increasing impact of that site on binding ability. Selection multipliers are chosen based on selection strength, *m*. Unless otherwise specified, *m* = 2 throughout this work. CDR positions each have a selection multiplier randomly drawn from an exponential distribution such that the multiplier is always greater than or equal to 1 and 99.5% of multipliers are less than *m*. FWR positions have a selection multiplier that is constructed similarly. A value is randomly drawn from an exponential distribution such that the value is always greater than or equal to 1, and 98% of values are less than *m*, then the square-root of the value is taken as the multiplier. By default, the distribution of site multipliers allows for most sites in the CDRs to evolve with near neutrality except for substitutions matching the target sequence, which are highly rewarded. The FWR distribution allows fewer sites to mutate with near neutrality than in the CDRs, while most sites will operate under purifying selection, reflecting that certain sites are more important to the structure and functionality of the FWR. The affinity score represents the square root of the product of the site multipliers for sites that match between the BCR and target AA sequence.

Let ***S*** be the set of positions that an amino acid sequence *A* has in common with the target sequence *T*, so that

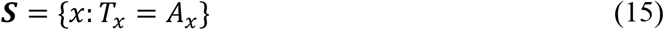

where *A* is the amino acid sequence, *A*_*x*_ is the *x*th position of *A, T* is the target AA sequence, and *T*_*x*_ is the *x*th position of *T*.

Then the affinity score *α* of the amino acid sequence *A* is:

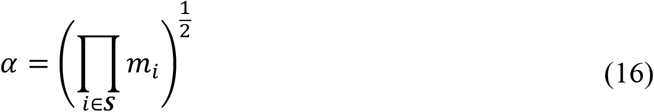

The affinity score of a cell is the sum of the affinity scores of its heavy chain and light chain. If a cell has a premature stop codon in either chain, its affinity is set to zero. Biologically, the affinity score of a cell represents how well that cell’s BCR binds to the target antigen. To simulate competition and affinity-driven selection, SimBLE GCs have a fixed number of antigen molecules available, which are distributed to cells with probability proportional to their relative affinity. Cells proliferate according to the number of antigen molecules bound, up to 10 child cells. Reproduction in the germinal center is modeled as sequential rounds of proliferative clonal bursting with transiently silenced SHM, followed by SHM of descendent cells.^44,45^ Algorithmically, this is equivalent to a cell having *c* direct children who each undergo one round of mutation. Thus, cells proliferate according to their amount of antigen bound by having *c* direct children, where *c* = min(10, number of antigen molecules bound). As such, if a cell binds to no antigen, it dies within a generation. Under neutral evolution, affinity is not calculated and antigen particles are distributed to cells randomly with equal probabilities rather than with probabilities weighted by affinity (**Supplemental Fig. 2**).

To model differentiation of MBCs and plasma cells, for each generation we draw *n*, the number of cells to differentiate and exit the GC, from a Poisson distribution with rate specified by the user (set to 2 cells/generation in all analyses presented here). Based on the current generation, *t*_*gen*_ and generations per day parameter (default 2 generations per day), SimBLE determines what percentage of *n* should become memory B cells (MBCs), *x*_*MBC*_, and what percentage should become plasma cells (PCs), *x*_*PC*_, according to observed kinetics of differentiation.^43^ From observed patterns, SimBLE sets

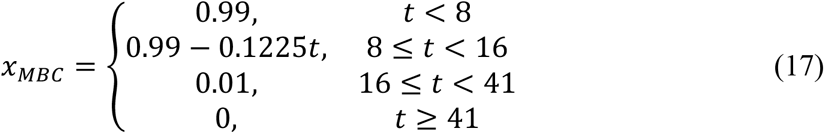

where *t* = *t*_*gen*_ · generations per day and represents the current day.

SimBLE then randomly selects *n*_*MBC*_ = *int*(*x*_*MBC*_ · *n*) cells from the GC population to differentiate into MBCs and selects *n*_*PC*_ = *n* − *n*_*MBC*_ cells to differentiate into PCs from the GC population with probability proportional to their affinity. This matches observed stochasticity of MBC differentiation, particularly in regard to affinity and high-affinity PC differentiation.^70,71^ The newly-differentiated MBCs and PCs are removed from the germinal center and added to the population in the “other” compartment.

### Simulating evolution of other sequence types using SimBLE

For applications not involving GC reactions, SimBLE can simulate uniform neutral evolution of random sequences from heterogeneously evolving populations. In this case, a randomly generated nucleotide sequence of user-specified length is used as the starting sequence rather than an observed naïve BCR, mutations are introduced randomly rather than according to known hotspot motifs^41^, and antigen is distributed to cells randomly with equal probabilities rather than with probabilities weighted by affinity (**Supplemental Fig. 2**).

### Pipeline for standard TyCHE analysis

To facilitate using TyCHE under similar specifications on multiple lineages, we developed a template-based system for running TyCHE from within the R package Dowser v2.4.1. BEAST2 uses XML files to specify data, model structure, prior values, and MCMC settings. We developed customizable XML templates which can be applied to multiple lineages at once. Users process data within the standard Dowser pipeline and can then specify the XML template with the desired model specifications using the getTimeTrees or getTimeTreesIterate functions. The function getTimeTrees will run BEAST2 with a specified XML template to a collection of lineages for a specified number of MCMC steps. Expanding on this function, getTimeTreesIterate will run getTimeTrees under the specified settings, assess convergence of desired parameters (e.g. effective sample size of at least 200), and resume the MCMC chain until either all parameters have converged or the maximum number of iterations is met. For analyses that include germline sequences (e.g. B cells and tumors), the rootfreqs package is used to set the MRCA as identical to the germline sequence.^72^ To allow for fixed differences between the germline and observed sequences, the germline sequence is also included as a tip whose date is set identical to the date of the root of the tree. Within Dowser, tree objects are imported using treeio and plotted using ggtree packages.^73,74^

### Simulation-based validation

TyCHE was validated through simulation-based analyses using simulations of primary and recall GCs under selective evolution and uniform neutral evolution. In all simulations, 20 clones were simulated and a total of 96 cells were sampled from GC and non-GC compartments at four distinct timepoints. In primary GC reaction simulations, 24 B cells are sampled at 50, 100, 150, and 200 generations. To simulate recall GC reactions, we simulated a primary GC for 100 generations, sampling 12 cells from the GC and 12 cells from the other compartment at generations 50 and 100. After the primary GC reaction finished, 24 non-GC B cells persist, from which one memory B cell was randomly selected to initiate a recall GC reaction. Functionally, the age of this cell was increased by 1000 generations, and we ran a new SimBLE simulation with this cell as the progenitor, beginning at generation 1100. The remaining 23 persistent non-GC B cells from the primary reaction were able to be sampled during the recall GC. To represent a new perturbation, such as a new vaccine, the target sequence for the secondary GC reaction was a mutated version of the original GC reaction’s target antigen, with 5 additional single nucleotide AA substitutions in the heavy chain and 3 single nucleotide AA substitutions in the light chain. Other than the mutated target sequence, the secondary GC reaction is configured to use identical model parameters and settings to the first GC reaction, sampling 12 B cells from the GC and 12 cells from the other compartment at generations 1150 and 1200.

TyCHE was benchmarked against the existing SC and UCLD models using standard discrete trait reconstruction.^30,39^ We sampled posterior distributions using MCMC with a chain length of 10^8^ iterations for primary GC simulations and 5 × 10^8^ iterations for recall GC simulations via the getTimeTreesIterate function from Dowser, with up to 9 additional chains of the same length or a maximum run time of one week for primary GC simulations and three weeks for recall GC simulations. While all models converged for primary and recall GC simulations under selection, while in uniform neutral simulations of recall GCs, 6 clones did not converge for TyCHE within the specified number of iterations. In primary GC reaction simulations, all models assumed GC-irreversibility, i.e., GC B cells were allowed to differentiate into non-GC B cells, but not the reverse. Naturally, in recall GC reaction simulations all models assume reversibility, i.e., GC B cells can differentiate into non-GC B cells and non-GC B cells can re-enter the GC.

For TyCHE, the GC rate of evolution *μ*_*GC*_ was initially estimated by the SC model applied to solely the GC B cells (only the primary GC reaction in the recall GC simulations). Then, the GC clock rate was given a strong normal prior distribution with mean equal to *μ*_*GC*_ and standard deviation equal to 10^−2^ × *μ*_*GC*_. The non-GC B cell clock rate was given a weaker normal prior distribution with mean 1 × 10^−6^ and standard deviation 1 × 10^−3^. For all models, we used a Bayesian Skyline tree prior with 5 intervals, the HKY substitution model,^68^ a lognormal prior on the transition/transversion rate ratio kappa with *M* = 0.67 and *S* = 0.2 (representing a mean ∼2), and empirical nucleotide frequencies. In simulated recall GC reactions, where GC to non-GC transitions were reversible, we set the root cell type to GC and the maximum tree height to 1000 generations above the true tree height unless specified otherwise (**Supplemental Fig. 3**).

The getTimeTreesIterate function in Dowser was used to sample the posterior distribution using MCMC with a chain length of 10^8^ steps and 2000 samples, with 200 labeled tree samples. If convergence was not reached, the MCMC chain was resumed for an additional 10^8^ steps up to 10 times. Convergence was evaluated with the criteria of all parameters operated on, except rate categories and internal node states, reaching an effective sample size of greater than 200.

Estimated time trees were compared to the ground truth from SimBLE. Specifically, we assessed the tree height (time from the most recent tip to the root), tree length (sum of all branch lengths), and the Robinson-Foulds distance after collapsing branches with length less than 5 generations for the evaluation of tree topology.

Differentiation times in simulated (**Fig. 3**) and empirical analyses (**Figs. 4 and 5**) were calculated using the height of each sampled non-GC B cell’s most recent ancestral GC node in the maximum clade credibility tree. In TyCHE, this height was reduced by the expected occupancy in the GC state of the intervening branch, which represents the amount of time in the GC between the parent GC node and descendant non-GC node. When subtracted from the age of the most recently sampled tip, this modified height indicates the estimated date of differentiation from a GC B cell. For non-TyCHE models differentiation dates were calculated similarly, but without the expected occupancy adjustment.

### Analysis of chronic GCs in HIV

Bulk BCR sequences from chronic GC reactions during HIV-1 infection were obtained from a prior study of three HIV-1 infected patients.^32^ Briefly, lymph node biopsies were obtained from donors H1, H2, and H3 (donors HIV1, 2, and 3 in original study, respectively), who were estimated to have been infected with HIV-1 for approximately 12-18mo, 24-36mo, and 4-8mo based on prior testing history and strength of anti-HIV antibody titers (Dr. Susan Moir, personal communication). Using FACS, B cells were sorted into GCBC, unswitched MBC, CD19^hi^ MBC, and CD19^lo^ MBC populations, which were sequenced separately using PCR-based amplification of BCR genes and Illumina MiSeq. Preprocessing, quality control, and identification of B cell clones were performed in ^32^.

Because sequences were collected from a single timepoint, it was not possible to estimate clock rates from these data. Instead, we estimated a clock rate using a prior longitudinal study of an HIV-1 infected patient over the first 3 years of infection.^47^ Measurably evolving lineages from this dataset, which accumulated a significantly higher SHM frequency over the sampled time period, were previously identified in ^58^. We estimated the clock rate of each of these lineages as the slope of its root-to-tip regression. We used the mean of these slopes (1 × 10^−3^ mut/site/week) as the mean clock rate for GCBCs, and an arbitrarily low clock rate of 1 × 10^−6^ for MBCs. These values were used as the means of strong normal prior distributions with sigma values of 10^−2^ × each clock rate, effectively fixing them. The overall rate of type switching was estimated using MCMC with an initial value of 1 switch/52 weeks. We used a Bayesian Skyline tree prior with 5 intervals as well as a lognormal prior on the transition/transversion rate ratio kappa with *M* = 0.67 and *S* = 0.2. For each relative transition rate, we used a gamma distribution with shape *α* = 0.1 and rate *β* = 1. As these were likely primary GC reactions, we assumed type switches from GC to non-GC were irreversible.

The getTimeTreesIterate function in Dowser was used to sample the posterior distribution via MCMC with a chain length of 10^8^ steps and 2000 samples, with 200 labeled tree samples. Convergence was assessed by whether all parameters that were operated on, except rate categories and internal node types, reached an effective sample size of at least 200. If convergence was not reached, the MCMC chain was resumed for an additional 10^8^ steps up to 15 total times. Clones that did not reach convergence were discarded. Clones were downsampled to represent GC and non-GC B cells as evenly as possible, with a maximum of 25 cells in each population per clone. Clones with fewer than 20 distinct BCR sequences were discarded.

### Analysis of repeated influenza vaccination

Single-cell RNA and BCR sequences were obtained from a prior study from three donors (P04, P05, P11) that received the 2018 and 2019 Flucelvax QIV from Sequirus.^33^ Because LN samples were not available from P11, only data from donors P04 and P05 were used in this study. No donors had received a flu vaccine in the preceding three years. All donors received the 2018 QIV at week 0. P04 received the 2019 QIV at week 35 and P05 received the 2019 QIV at week 38. Samples were obtained from peripheral blood mononuclear cells (PBMCs), lymph node fine needle aspirations (FNAs), and bone marrow plasma cells (BMPCs). Samples from P04 were obtained from sorted plasmablasts (PBs) at weeks 1 and 36, enriched IgD^lo^ B cells at weeks 0, 2, 17, 35, 37, 48, whole FNA at weeks 0, 1, 2, 17, 35, 37, 44, 48, and enriched BMPCs at weeks 0, 4, 26, 35. For P05, samples were obtained from whole PBMCs at weeks 13 and 17, enriched IgD^lo^ B cells at weeks 13, 17, 26, 38, 39, 40, 42, 47, 51, whole FNA at weeks 13, 26, 38, 39, 42, 47, 55, and enriched BMPCs at weeks 0, 4, 13, 26, 38, 42, 51. Samples were sequenced using the 10X Genomics Single Cell 5′ RNAseq with BCR amplification. Preprocessing, quality control, and identification of B cell clones and subtypes were performed in.^33^ To determine whether particular clones reacted to influenza, monoclonal antibodies (mAbs) were previously derived from selected clones and tested for binding affinity to influenza antigens. Clones were determined to be measurably evolving if they accumulated significantly higher SHM frequency over the sampled time period as determined by the correlationTest function in Dowser.^58^ We analyzed all clones containing an influenza-binding mAb sequence, that had GC B cells sampled after both vaccinations, and were measurably evolving. This resulted in 1 clone from P04 and 7 from P05. Clones with greater than 100 total sequences were down-sampled to 100 sequences while ensuring maximum balance between sampled B cell subtypes and tissues.

For type-linked analysis with TyCHE, the GC clock rate was given a normal prior distribution with mean equal to 4.9 × 10^−3^ mut/site/week and standard deviation equal to 10^−2^ × the mean. The clock rate for the other B cell subtypes was given a normal prior distribution with mean equal to 1 × 10^−6^ and standard deviation equal to 1 × 10^−3^. The GC clock rate was estimated as the median slope of the root-to-tip regression for previously identified measurably evolving lineages from P05 sampled after the 2018 QIV vaccination (days 0-60).^48,58^ For all models, we used a Bayesian Skyline tree prior with 5 intervals and a lognormal prior on the transition/transversion rate ratio kappa with an *M* = 0.67 and *S* = 0.2. The priors on all cell type relative transition rates were set to gamma distributions with shape = 0.1 and scale = 1.0. To help with convergence and identifiability in this reversible scenario (**Supplemental Fig. 3**), the cell type at the root was fixed to GC and the maximum root date of the trees was set to four years prior to the 2018 QIV vaccination, as neither patient had received a flu vaccine in the preceding three years. While the actual root date of these lineages was potentially before 2014, our ability to resolve this date was low due to the arbitrarily low clock rate of MBCs. Thus, estimated root dates may be more recent than their true values.

The getTimeTreesIterate function in Dowser was used to sample the posterior distribution via MCMC with a chain length of 10^9^ steps and 2000 samples as well as 200 state-labeled tree samples. If convergence was not reached, the MCMC chain was resumed for an additional 10^9^ iterations up to 10 total times. Sampled posteriors were determined to have converged if all parameters that were operated on, excepting rate categories and internal node types, reached an effective sample size of at least 200.

When determining the proportion of branches predicted to be GC B cells over time for TyCHE (**Fig. 5B, Supplemental Figs. 5-6**), we used the expected occupancy in the GC state estimated by the model for that branch. For non-TyCHE models, we assumed that all branches are in the same state as the child node. For determining GC reaction start dates in **Fig. 5D**, we defined a GC reaction as a contiguous set of internal nodes predicted to be GC B cells that lead to sampled GC B cells in the maximum clade credibility tree. We estimated the date that each GC reaction began as the date of its earliest GC internal node.

### Analysis of hypermutating glioma tumors

Sequences of glioma tumors were obtained from the GLASS Consortium.^34,75^ Patient 4F0A was selected due to multiple non-hypermutating samples and the presence of a hypermutator phenotype in one sample. Sequences from this patient included normal genome (NG), primary tumor (TP, day 0), first recurrence (R1, non-hypermutator based on mutation level, day 3042), and second recurrence (R2, hypermutator, day 3722). Clinical and sequencing records for patient case barcoded GLSS-HF-4F0A were downloaded from the Synapse Repository (syn17038081). Variant calls from the 2022 updated consensus variant call format (VCF) files originally published in ^34^ were additionally filtered for likely germline variants by comparing them to recent releases of dbSNP (2025-03-12) and gnomAD (2024-10-11). Multiple sequence alignments were generated by first filtering VCF files to variants of size 1. Then, for any site mutated in at least one of the sequenced samples, a concatenated alignment was generated across mutant sites, using the paired-normal reference allele in instances where a mutation was not observed in a particular sample.

Genetic distance tree topologies, branch lengths, and model parameters were estimated using the *dnaml* program of PHYLIP v3.695.^76^ We estimated the mean clock rate of non-hypermutators (N) using a strict clock model applied to only NG, TP, and R1. Due to the limited number of sequences, we estimated the clock rate of the hypermutator (H) using a root-to-tip regression from the genetic distance tree of NG, R1 and R2. For cell type specific clock rates in TyCHE, we used a normal prior with a mean equal to the estimated clock rates of H and N strains and sigma values equal to their respective mean × 10^−3^, effectively fixing these rates. Switches were assumed to be irreversible from N to H. Due to low sequence count, all models used a coalescent constant population size tree prior. A prior on kappa was set using a lognormal distribution with *M* = 1.25 and *S* = 0.5. Empirical nucleotide frequencies were used. The getTimeTreesIterate function in Dowser was used to sample the posterior distribution via MCMC with a chain length of 10^9^ iterations and 2000 samples and 200 state-labeled tree samples. Convergence was assessed by whether each parameter operated on, except rate categories and cell type states, reached an effective sample size of at least 200. If convergence was not reached, the MCMC chain was resumed for an additional 10^9^ iterations up to 20 times.

### Analysis of P. aeruginosa lung infection

*P. aeruginosa* sequence alignments were obtained from a prior study.^25^ Isolates were obtained from bronchial alveolar lavage samples from a patient admitted for COVID-19 related pneumonia with no sign of *P. aeruginosa* lung infection at 46, 48, 76, 81, 83, 89, 92, and 101 days after admission. Isolates were sequenced using Illumina NextSeq. Sequence processing, assembly, SNP identification, and sequence alignment were performed as detailed in.^25^ For computational expediency, only sites with SNPs were included in the alignment. Genetic distance tree topology, branch lengths, and HKY model parameters were estimated using maximum likelihood as implemented in the *phangorn* R package.^77^ Clock rates and tree heights of the non-hypermutator (N) and hypermutator (H) strains were estimated using SC models on those sequences separately. For TyCHE models, we used these mean clock rates as the mean of the normal prior for clock rates of N and H respectively, with a sigma value of 10^−3^ × the prior mean. This effectively fixed the clock rate for each type. Switches from N to H were assumed to be irreversible. In all models, the HKY substitution model was used as well as a Bayesian Skyline tree prior with 5 intervals. A prior on the transition/transversion rate ratio was set using a lognormal distribution with *M* = 1.25 and *S* = 0.5. The getTimeTreesIterate function in Dowser was used to sample the posterior distribution via MCMC with a chain length of 10^9^ iterations and 2000 samples Convergence was assessed by whether all parameters that were operated on, except rate categories and internal node types, reached an effective sample size of at least 200. If convergence was not reached, the MCMC chain was resumed for an additional 10^9^ iterations up to 10 times.

## Notes

### Competing Interest Statement

The authors have declared no competing interest.

### Summary of Updates

Improved underlying models and functionality of SimBLE and TyCHE. Updated Figs 3-5 to include differentiation timing applications. Updated author list.

