## Supplemental Information for "TyCHE enables time-resolved lineage tracing of heterogeneously-evolving populations"

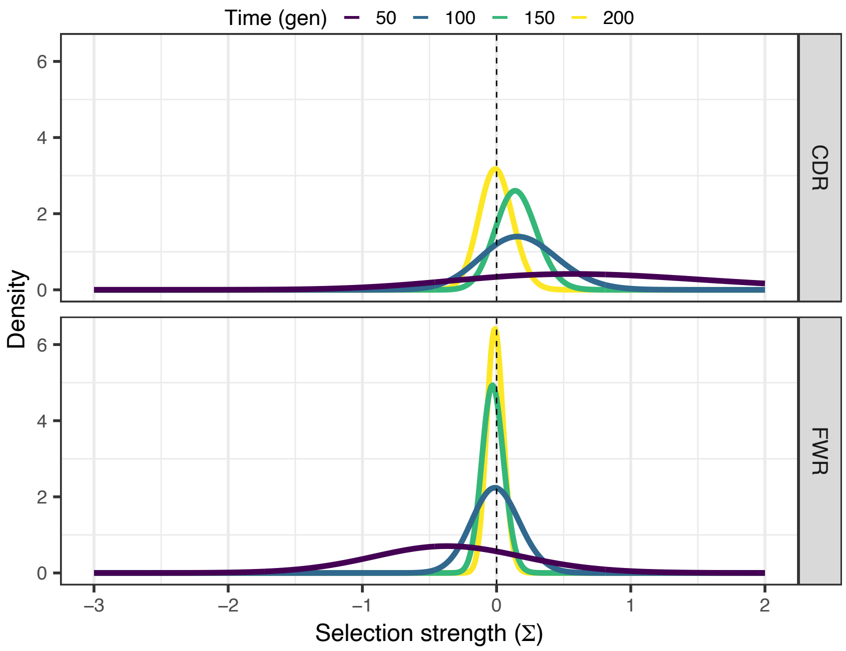


**Supplemental Figure 1:** BASELINe analysis of neutral simulations. Posterior distributions of selection scores from BASELINe applied to BCRs from 100 lineages sampled at 50, 100, 150, and 200 generations under simulations without selection. 5653 sequences with in-frame stop codons were removed as BASELINe cannot determine selection scores when consensus sequences contain early stop codons. Sequences sampled at each generation were analyzed separately. Values below zero indicate purifying selection while values greater than zero indicate diversifying selection. CDR indicates antigen-binding complementarity-determining regions while FWR indicates structural framework regions.

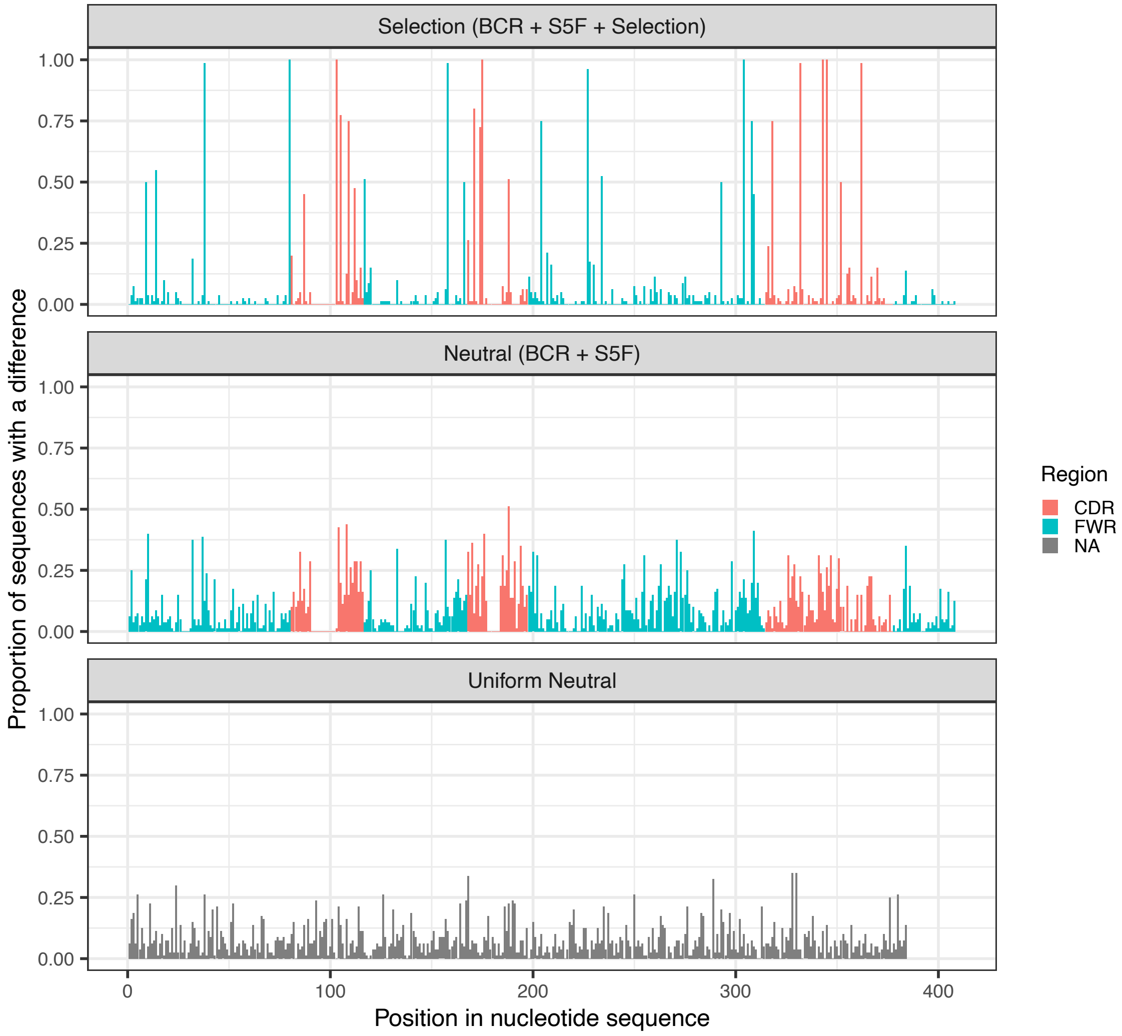


**Supplemental Figure 2:** Proportion of sequences with a difference at each nucleotide position for 1 clone under selection (starting BCR sequence + S5F mutation and substitution model + selection), 1 clone under BCR neutral simulation (starting BCR seq + S5F mutation and substitution model), and 1 clone under uniform neutral simulation (random starting sequence + uniform substitution probabilities). Neutral and selection models use IMGT gapped sequence position and positions are colored by region (CDR, FWR). Uniform neutral sequences have length 384 and BCR neutral and selection have un-gapped sequences of length 384.

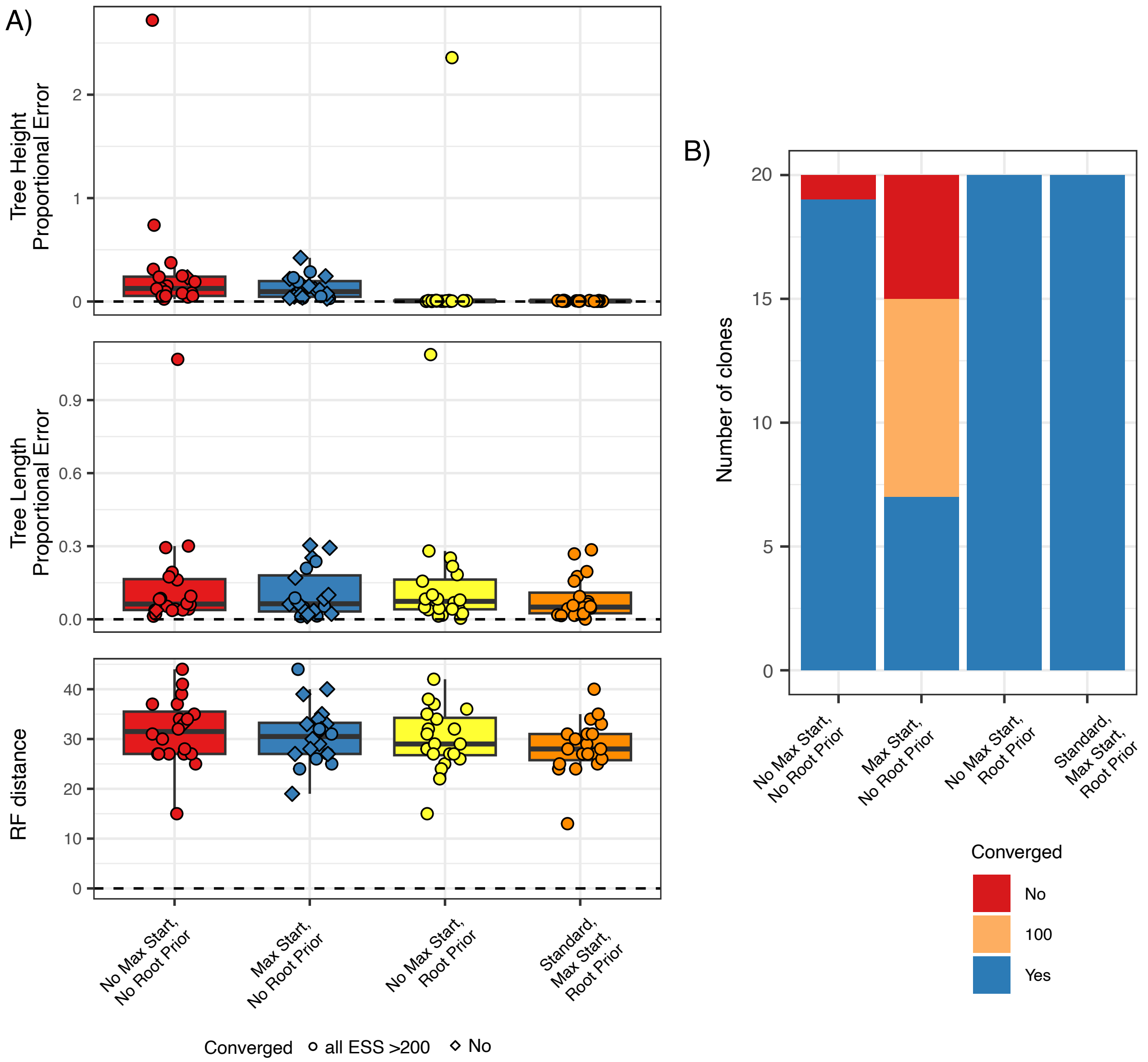


**Supplemental Figure 3:** Benchmarking of TyCHE with and without constraints, for recall GC simulations. For all configurations with "root prior", the root cell type was fixed to GC, and with "max start", the maximum tree height was set to 1000 generations (~83%) above the true tree height. The "standard" configuration which was used throughout our other analyses uses both constraints. We sampled posterior distributions using MCMC with a chain length of $5x10^8$ iterations via Dowser's getTimeTreesIterate function, with up to 9 additional chains of the same length or a maximum run time of two weeks. **A)** Each dot represents one simulated clone. From top to bottom: proportional error of estimated tree height (generations from root to most recent tip), dashed line at 0 represents no error; proportional error of tree length estimates, calculated as the sum of all branch lengths, dashed line at 0 represents no error; Robinson-Foulds distance from the true tree topology, dashed line at 0 represents identical topology to the true tree. To reduce noise, branches under 5 generations were collapsed for Robinson-Foulds distance calculation. **B)** Summary of convergence after maximum run time (14 days). "Yes" represents clones for which all parameters had an ESS > 200, "100" represents clones for which all parameters had an ESS > 100, and "No" represents clones which had any parameters with lower ESS values.

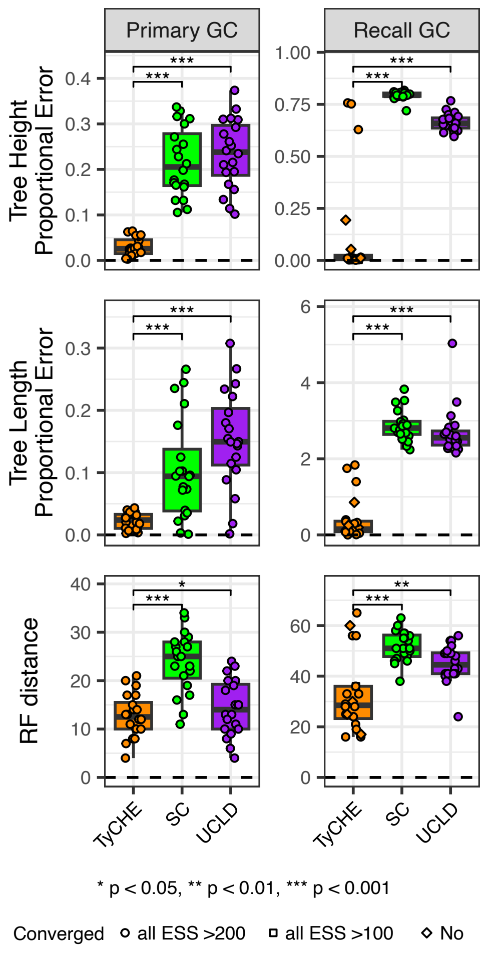


**Supplemental Figure 4:** Benchmarking of TyCHE and existing models on uniform neutral simulations, for primary (left) and recall (right) GC simulations. Each dot represents one simulated clone. From top to bottom: proportional error of estimated tree height (generations from root to most recent tip), dashed line at 0 represents no error; proportional error of tree length estimates, calculated as the sum of all branch lengths, dashed line at 0 represents no error; Robinson-Foulds distance from the true tree topology, dashed line at 0 represents identical topology to the true tree. To reduce noise, branches under 5 generations were collapsed for Robinson-Foulds distance calculation. P values were calculated using the paired Wilcoxon signed-rank test.

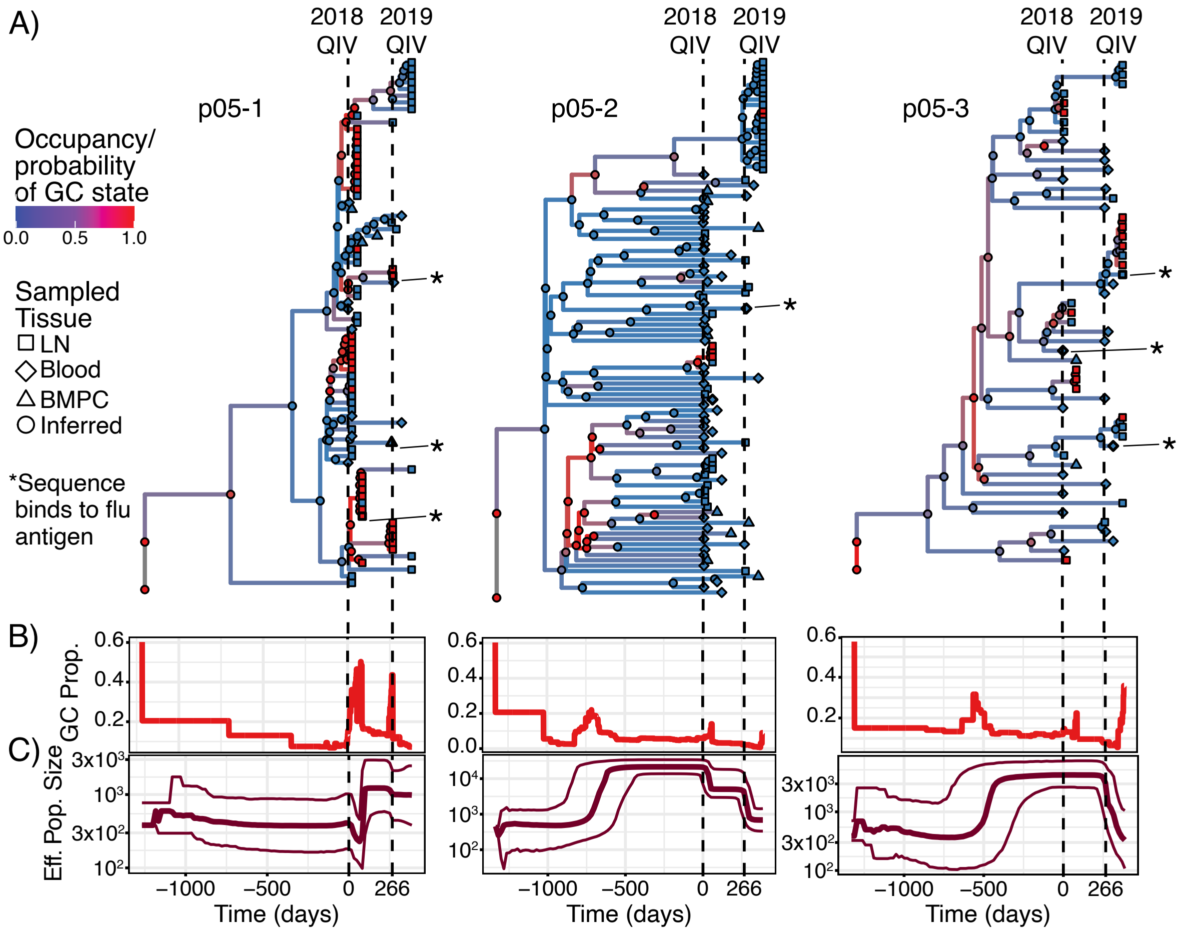


**Supplemental Figure 5:** Time-resolved analysis of recall GC reactions following repeated influenza vaccination of donor P05 (clones 1-3). The donor was immunized with 2018/2019 QIV vaccination at week 0, and re-immunized with 2019/2020 QIV at week 38. Samples were taken from PBMCs, lymph node (LN) fine-needle aspiration (FNA), and bone marrow plasma cells (BMPCs). **A)** TyCHE-inferred time trees. Sequences experimentally confirmed to bind influenza antigens are marked with *. Dashed lines indicate dates of influenza vaccination. **B)** Proportion of branches predicted to be GC B cells over time. **C)** Bayesian skyline plots showing changes in effective population size.

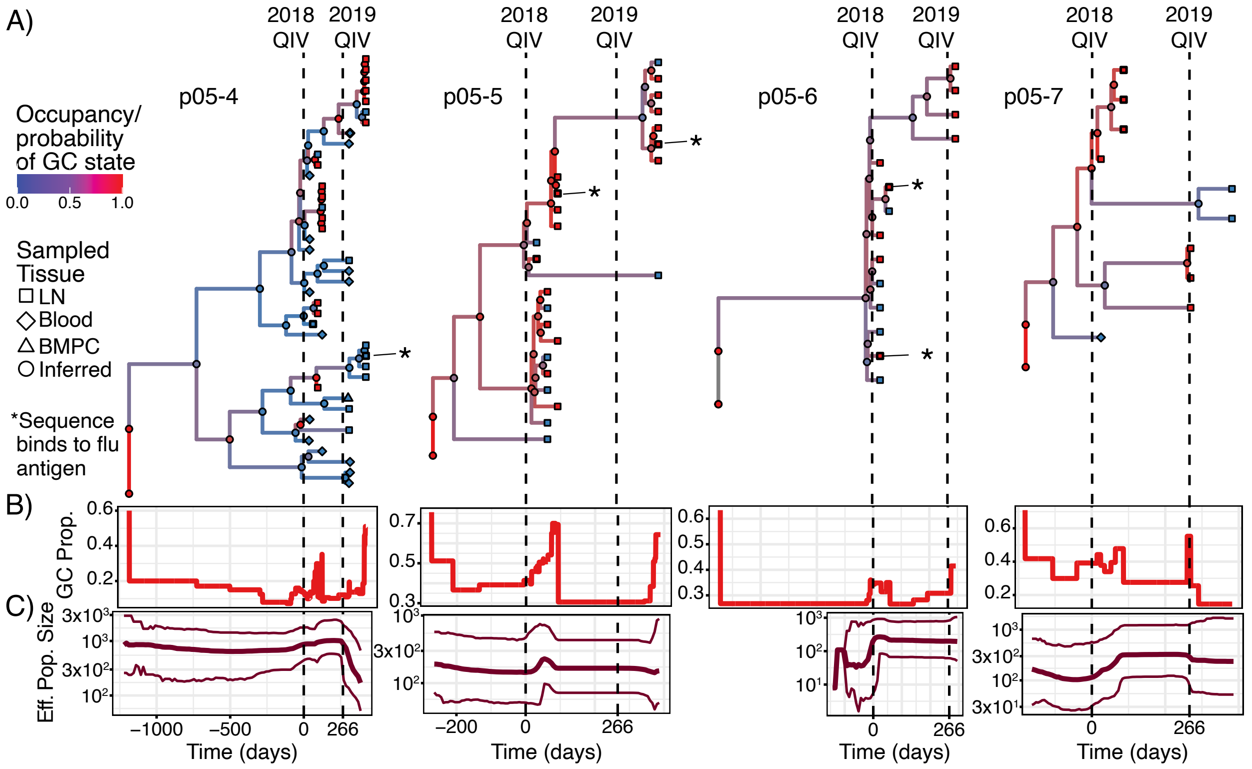


**Supplemental Figure 6:** Time-resolved analysis of recall GC reactions following repeated influenza vaccination of donor P05 (clones 4-7). The donor was immunized with 2018/2019 QIV vaccination at week 0, and re-immunized with 2019/2020 QIV at week 38. Samples were taken from PBMCs, lymph node (LN) fine-needle aspiration (FNA), and bone marrow plasma cells (BMPCs). **A)** TyCHE-inferred time trees. Sequences experimentally confirmed to bind influenza antigens are marked with *. Dashed lines indicate dates of influenza vaccination. **B)** Proportion of branches predicted to be GC B cells over time. **C)** Bayesian skyline plots showing changes in effective population size.

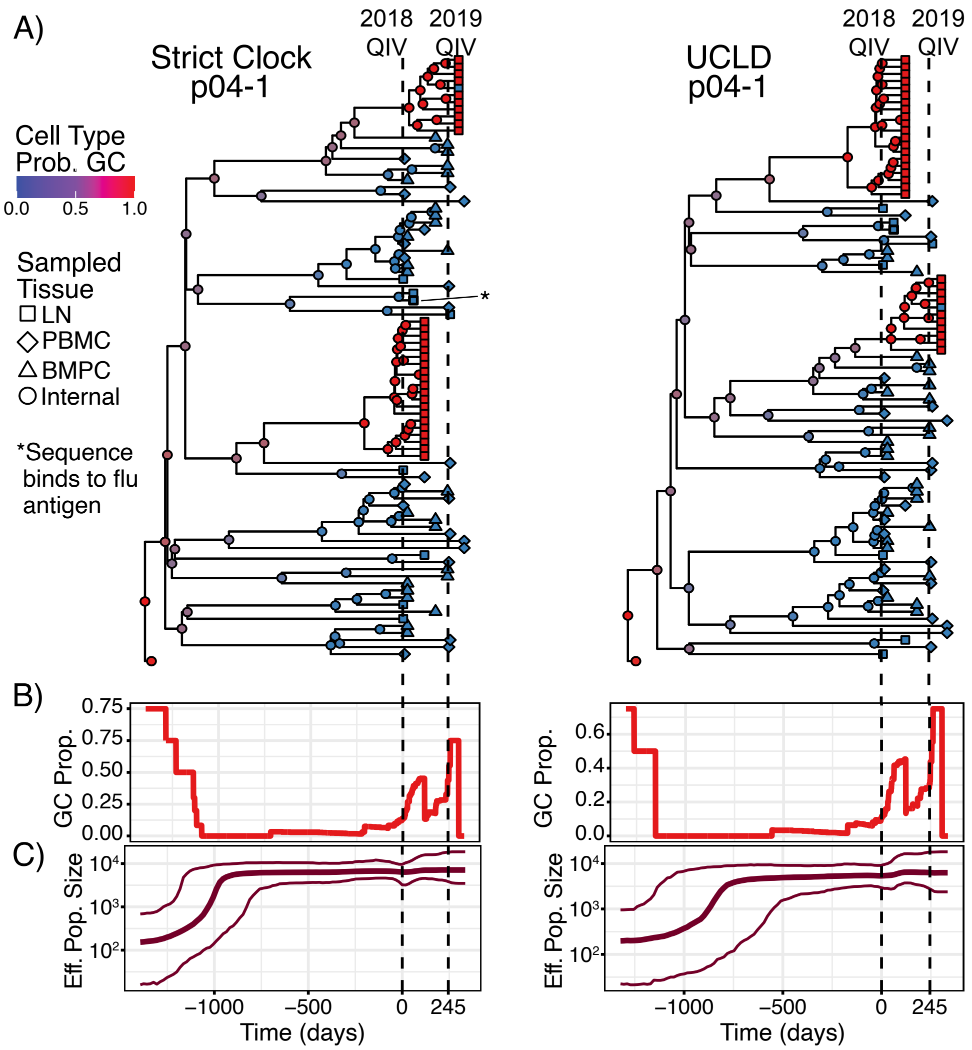


**Supplemental Figure 7:** Strict Clock (SC) and UCLD time-resolved analysis of recall GC reactions following repeated influenza vaccination of donor P04, clone 1. This clone is shown in Fig 5 A-C. The donor was immunized with 2018/2019 QIV vaccination at week 0, and re-immunized with 2019/2020 QIV at week 35. Samples were taken from PBMCs, lymph node (LN) fine-needle aspiration (FNA), and bone marrow plasma cells (BMPCs). **A)** SC- and UCLD-estimated time trees. Sequences experimentally confirmed to bind influenza antigens are marked with *. Dashed lines indicate dates of influenza vaccination. **B)** Proportion of branches predicted to be GC B cells over time. **C)** Bayesian skyline plots showing changes in effective population size.
